# A bacterial Rhesus transporter retunes a structurally conserved ammonium pore into a reversible nitrogen valve

**DOI:** 10.64898/2026.09.24.754001

**Authors:** Adriana Bizior, Mingyu Qin, Peter Henderson, Benjamin Cooper, Daniel E. Larcombe, Paul A. Hoskisson, Georgia Isom, Leighton Pritchard, Syma Khalid, Arnaud Javelle

## Abstract

How conserved proteins acquire new physiological functions is a central question in molecular evolution. Rather than inventing new architectures, evolution often repurposes existing scaffolds, preserving core structural features while retuning the molecular logic that connects mechanism to physiology. Membrane transporters offer a powerful test of this principle because substrate selectivity, directionality, flux and energetic cost must be coordinated at the interface between the cell and its environment.

The Amt/Mep/Rh superfamily controls one of the most fundamental requirements of cellular life: the movement of reduced nitrogen across biological membranes. Despite sharing a highly conserved ammonium-conducting pore, these proteins support distinct physiological roles, including nitrogen acquisition, sensing and homeostatic control. Here, we combined targeted mutagenesis, electrophysiology, yeast complementation and molecular dynamics simulations to define the transport logic of NeRh50, a bacterial Rhesus protein from the ammonia-oxidising bacterium *Nitrosomonas europaea*. We show that NeRh50 is not simply an AmtB-like ammonium importer. Instead, it uses the conserved Amt/Mep/Rh pore as a branched transport system in which inward uptake and export-linked transport can be separated genetically and mechanistically. Two conserved pore residues define this division of labour. A residue at the external entrance couples ammonium recruitment to productive inward uptake, whereas a second residue deeper in the pore enables a distinct transport mode required for substrate release when intracellular nitrogen accumulates. Thus, conserved Amt/Mep/Rh pore landmarks do not impose a single mechanism. Their local chemistry can be reassigned to generate different transport outputs, allowing NeRh50 to function as a reversible nitrogen valve. These findings reveal how minimal retuning within an ancient membrane-protein scaffold can rewire transport directionality and adapt nitrogen handling to ecological and physiological demand.

## Introduction

New physiological functions often emerge during evolution through modification of existing protein scaffolds rather than through the invention of entirely new ones. Enzyme superfamilies provide a classic example, in which conserved folds retain a common chemical logic while enabling distinct metabolic reactions^1,2^. Membrane transporters extend this principle to the interface between the cell and its changing environment, where substrate specificity, directionality, transport flux and energetic cost must be coordinated with cellular physiology and adapted to an ecological niche^3–5^. The ammonium transporters of the Amt/Mep/Rh superfamily control a universal requirement of cellular life: the movement of reduced nitrogen across biological membranes. This process is essential for biosynthesis, energy metabolism and nitrogen homeostasis, yet intracellular ammonia/ammonium accumulation is toxic. The intrigue of the Amt/Mep/Rh superfamily is that one highly conserved ammonium pore can be modified to support distinct physiological outputs, from nitrogen acquisition and sensing to homeostatic control. The family therefore provides a particularly informative system for understanding how evolution retunes a conserved membrane scaffold to meet different physiological demands^6–8^. Crystal structures of *Escherichia coli* AmtB established the canonical architecture of the transporters, with each monomer containing a narrow hydrophobic pore, an extracellular ammonium-recruitment site and a twin-His motif within the conducting pathway^9,10^. Subsequent structures of archaeal Amt, fungal Mep and eukaryotic Rh proteins confirmed conservation of this transmembrane core across the superfamily^11–16^. The unresolved question is therefore not whether Amt/Mep/Rh proteins share an architecture, but how that conserved architecture has been retuned to produce distinct transport logic: Amt/Mep proteins are generally associated with inward ammonium acquisition and, in some cases, signalling, whereas Rh proteins have been linked to ammonia/ammonium homeostasis and bidirectional or protective ammonium handling^17,18^. Mechanistic studies have identified conserved pore residues, including a pore-entry aspartate and the twin-His motif within the hydrophobic pore, as key determinants of ammonium translocation^13,19–22^. However, most of this work has examined individual proteins in separate experimental systems. It therefore remains unclear whether conserved residues perform equivalent functions in Amt and Rh proteins, or whether their roles have diverged during evolution. Because these residues are central to current models of ammonium transport^20,23,24^, resolving this question requires direct comparison of Amt-type and Rh-type transport in the same experimental framework.

*Nitrosomonas europaea* provides a powerful system to address this problem. This obligate ammonia-oxidising bacterium depends on ammonia as both an energy substrate and a nitrogen source^25,26^, yet encodes no canonical Amt-type transporter and instead carries NeRh50, a bacterial Rh-family protein proposed to have been acquired by horizontal gene transfer from a eukaryotic lineage^27^. Recent structural and functional characterisation further showed that NeRh50 combines a distinctive lipid-associated architecture with selective, low-affinity electrogenic ammonium transport^16^. NeRh50 therefore offers a rare opportunity to test how an Rh protein controls ammonium flux in an organism whose physiology is built around ammonia metabolism, and to compare this mechanism directly with AmtB, the best-characterised paradigm member of the superfamily.

Here, using NeRh50 and AmtB as paired representatives of the Rh and Amt branches, we combine electrophysiology, solvent exchange, mutagenesis, yeast functional assays and molecular dynamics simulations to test how a conserved ammonium-transport scaffold has been functionally redeployed. We show that NeRh50 contains an Rh-like electrogenic transport state that is mechanistically distinct from canonical AmtB, and that inward ammonium uptake can be genetically uncoupled from its export-associated/protective activity. These findings reveal that conserved pore residues are not functionally equivalent across the Amt/Mep/Rh superfamily but have been evolutionarily repurposed to separate import from Rh-like protective/export-associated transport within the same molecular scaffold.

## Materials and Methods

### Bacterial strains, yeast strains and plasmids

*N. europaea* NeRh50 and *E. coli* AmtB constructs were expressed either for biochemical/electrophysiological analysis or for yeast functional assays. For protein purification and Solid Supported Membrane Electrophysiology (SSME) measurements, NeRh50 variants were expressed in *E. coli* GT1000 (ΔglnK, ΔamtB)^28^ from the pAD7 expression vector^27^. For yeast assays, NeRh50, *E. coli* AmtB and the corresponding point mutants were cloned into pDR195 under control of the PMA1 promoter^29^. Yeast complementation assays were performed in the ammonium-transporter-deficient *S. cerevisiae* strain 31019b (MATa ura3 mep1Δ mep2Δ::LEU2 mep3Δ::KanMX2)^30^. Methylammonium (MeA)-protection assays were also performed in a wild-type yeast background, strain 23344c, where endogenous Mep-dependent MeA entry can be modulated by nitrogen source^30^.

### Site-directed mutagenesis

Point mutations were introduced into NeRh50 or AmtB expression constructs by site-directed mutagenesis using primers carrying the desired substitutions. The NeRh50 variants analysed were D162A, H170A, H170D and H170E. Equivalent AmtB substitutions included D160A and H168A/D/E^20,31^. All constructs were confirmed by Sanger sequencing across the full coding region or the mutated region before use in protein expression or yeast assays.

### Protein expression and purification

NeRh50 and NeRh50 variants were heterologously overexpressed and purified as previously described^14^

### Proteoliposome preparation

Proteoliposomes were prepared from *E. coli* polar lipids and POPC mixed at a 2:1 ratio by weight as previously described^32^

### Solid-supported membrane electrophysiology

Solid-supported membrane electrophysiology (SSME) was performed using a SURFE^2^R N1 instrument (Nanion Technologies, Munich, Germany) as previously described^20,32^

### Yeast assay

All yeast assays were performed after two independent transformations, with each transformation assayed twice. AmtB D160A and AmtB H168 variants have been characterised previously in yeast-based assays^20,31^. However, those studies used AmtB-focused assay designs and different technical conditions, including different ammonium ranges, MeA concentrations and/or expression contexts. Here, the AmtB controls were repeated in parallel with NeRh50 and the corresponding NeRh50 variants using the same vector background, yeast strain, growth medium, nitrogen-source conditions and MeA concentrations. This allowed direct comparison of AmtB-like import behaviour and NeRh50-associated MeA protection/export activity within a single experimental framework.

### Ammonium-complementation assays

*NeRh50, amtB* and mutant constructs in pDR195 were transformed into the *S. cerevisiae* triple-mepΔ strain 31019b. Transformants were selected on YNB minimal medium lacking uracil. For complementation assays, cells were grown in buffered YNB medium at pH 6.1 containing 3% glucose and the indicated nitrogen source. Cultures were adjusted to OD600 0.6, washed three times in HEPES buffer pH 6.1, diluted 1:100 and 50 μL was plated onto selective medium. Growth was assessed on glutamate as a non-selective nitrogen-source control and on limiting ammonium concentrations, typically 0.5, 1 and 3 mM NH_4_ ^+^, as the sole nitrogen source. Plates were incubated at 30°C and imaged after 3 and 4 days.

### Methylammonium-protection assays in triple-mepΔ yeast

To assess MeA resistance in the absence of endogenous ammonium transporters, triple-mepΔ yeast expressing empty vector, WT AmtB, WT NeRh50 or the corresponding variants were plated on glutamate medium supplemented with methylammonium. MeA concentrations included 30, 100 and 200 mM, with the highest concentration used as the most discriminatory condition for NeRh50-associated protection. Growth in the presence of MeA was compared with empty vector, WT AmtB and WT NeRh50 controls.

### Methylammonium-toxicity assays in wild-type yeast

To examine MeA protection in a background where endogenous Mep transporters contribute to MeA entry, NeRh50 and AmtB constructs were also expressed in wild-type yeast. Nitrogen sources were selected to modulate endogenous Mep expression: proline and glutamate were used as Mep-permissive conditions, glutamine was used as a repressing nitrogen source, and glutamine supplemented with ammonium was used as a restrictive control combining Mep repression with ammonium competition. Cells were grown in the corresponding YNB medium supplemented with proline, glutamate, glutamine, or glutamine plus ammonium, then plated onto the same YNB/nitrogen-source medium in the absence or presence of MeA, as described above.

### Plate imaging and phenotype scoring

Yeast plates were imaged after 3 and 4 days of incubation at 30°C. Images were acquired as greyscale JPEG files. For display, a linear contrast stretch was applied identically to all images by subtracting the 20th intensity percentile as the agar baseline and rescaling so that the 98th percentile mapped to maximum intensity. Phenotypes were assigned by comparison with empty-vector, WT AmtB and WT NeRh50 controls under the same nitrogen and MeA conditions.

### Statistical analysis and dataset-specific controls and comparability across independent SSME experimental series

All SSME measurements were generated in two independent experimental series, each comprising purifications and reconstitutions performed with matched lipid, detergent and Bio-Beads batches. The two series were performed months apart using independent reagent batches, providing independent experimental replication. Series 1 characterised WT NeRh50 and pore variants in H_2_O at Lipid-to-Protein Ratio (LPR) 5 and LPR10, and these data were incorporated into the cross-series comparison in Figure 1, the WT/D162A LPR analysis in Figure 3C-D, and the H170 variant LPR analysis in Figure 5B. Series 2 examined paired H_2_O/D_2_O/recovery responses and the WT LPR series at LPR5, LPR10 and LPR50; these data were incorporated into the cross-series comparison in Figure 1, the WT solvent-exchange and LPR analyses in Figure 2B, D and F, the D162A solvent-exchange analysis in Figure 3F, and the H170 solvent-exchange analysis in Figure 5C.

SSME maximum amplitudes and decay rate constants were fitted separately using linear models with the relevant explanatory variables: Value ∼ Series for Figure 1, fitted independently for each genotype with heteroskedasticity-consistent standard errors (HC3, sandwich); Value ∼ LPR for Figure 2B and Figure 2F; Value ∼ Condition for Figure 2D, Figure 3F and Figure 5C; and Value ∼ LPR × Variant for Figure 3C-D and Figure 5B. Models were fitted in R v4.4.3^33^ using the lm() function, with model parameters extracted using the parameters package v0.28.3^34^, model assumptions assessed using the performance package v0.16.0^35^, and pairwise contrasts estimated using emmeans v2.0.2^36^. Concentration-response data were fitted to the Michaelis-Menten equation by nonlinear regression to estimate apparent K_m_ values using GraphPad Prism version 11.0.0 for Windows^37^.

### Molecular dynamics simulations

#### System preparation

Simulations were based on the NeRh50 structure^14^. The protein was embedded with CHARMM-GUI Membrane Builder in a POPE:POPG:cardiolipin bilayer at an 80:15:5 molar ratio^38^. The system was solvated with CHARMM-modified TIP3P water, neutralised and adjusted to 0.15 M NaCl in a box of approximately 15 × 15 × 12 nm^3^. Protein, lipids and ions were described with the CHARMM36m force field^39^; NH_4_ ^+^ used CHARMM ammonium atom types equivalent to the lysine NZ–HZ group.

#### Equilibration and production

Each system was energy minimised, equilibrated for 5 ns in the NVT ensemble, and propagated for 500 ns of unrestrained production in the NPT ensemble, with three independent replicates per system. Equations of motion were integrated with the leap-frog algorithm and a 2 fs timestep, and bonds to hydrogen were constrained with LINCS (order 4, warning angle 90°). Neighbour searching used the Verlet scheme on a grid with the pair list rebuilt every 20 steps (buffer tolerance 0.005). Electrostatics were treated with PME and a 1.2 nm real-space cutoff. Van der Waals interactions were force-switched between 1.0 and 1.2 nm, with the pair-list cutoff at 1.2 nm. Temperature was maintained at 310 K using velocity rescaling (τ_T = 1.0 ps), with protein, bilayer, and solvent coupled to separate baths. Pressure was held at 1 bar with the Parrinello-Rahman barostat applied semi-isotropically (τ_P = 4.0 ps, compressibility 4.5 × 10^-5^ bar^-1^). Simulations were performed with GROMACS 2021.5^40^.

#### Ammonium-loaded and electric-field simulations

Starting configurations were taken from the endpoints of the 500-ns production trajectories. Three NH_4_^+^ ions were introduced on the periplasmic side by replacing periplasm-facing Na^+^ ions near the pore mouth, one ion per protomer. For H170D, the Na^+^ count was adjusted to maintain charge neutrality after the three additional negative charges were introduced in the trimer. Each loaded system was simulated for 250 ns in three independent replicates. Field-driven runs used the same setup with a static field of 0.05 V nm^-1^ applied along the negative z-axis, parallel to the membrane normal and channel axis. Ion-resolved metrics were normalised per ion.

#### Umbrella sampling and potential-of-mean-force reconstruction

The reaction coordinate was the substrate position along the channel z-axis. Starting configurations were generated by steering the substrate through one WT protomer at 0.5 nm ns^-1^ with a force constant of 500 kJ mol^-1^ nm^-2^. Eighty-eight windows were placed at 0.05-nm intervals across 4.4 nm. Each window used a 1000 kJ mol^-1^ nm^-2^ harmonic restraint, was equilibrated for 5 ns and was sampled for 20 ns under the production thermostat and barostat. No additional protein restraints were applied. Potentials of mean force were reconstructed with the weighted histogram analysis method implemented in g_wham^41^. Calculations were performed separately for fixed-state NH_4_ ^+^ and NH_3_.

#### Trajectory coordinates and residue numbering

Periodic-boundary artefacts were corrected with gmx trjconv (-pbc mol -center), and trajectories were analysed with MDAnalysis^42^. Each protein was superposed on the Cα atoms of the first frame of the corresponding WT or H170D system. For trajectory analyses, z was measured relative to the trimer centre of mass and positive values were periplasmic. Per-protomer axes joined the Cα centroids of the Phe-gate and twin-His site and were recalculated in every frame. The simulation model is offset by 31 residues from the experimental precursor sequence: D131, F79, F187, H139 and H293 in the trajectories correspond to experimental D162, F110, F218, H170 and H324, respectively. Experimental numbering is used in the Results and figures. The PMF plots use the opposite orientation, with negative z values periplasmic; this panel-specific convention is stated in the Figure 8 legend.

#### Density, occupancy, hydration and pore-geometry analyses

Time-resolved axial NH_4_ ^+^ density maps were constructed by histogramming ion positions in time and z across the three replicates. Counts were divided by the numbers of ions and contributing frames. Three-dimensional densities were computed on a 1-Å cubic grid spanning the trimer, pooled across replicates and normalised to the estimated bulk ammonium concentration. A common isovalue, defined as a fixed fraction of the global maximum, was used for comparison. Per-residue occupancy was the probability that a residue lay within 4 Å of an ammonium nitrogen, normalised per ion. Axial wet fraction was calculated from water occupancy in each axial bin of the substrate-free trajectories. Effective pore-radius profiles were evaluated along each per-protomer axis, and gate geometry was assessed at the twin-His and Phe-gate constrictions. Profiles and distributions were pooled across the three independent replicates.

#### Position-dependent pK_a_

A continuous pK_a_ profile along the pore axis was derived from the two PMFs via a thermodynamic cycle. The local pK_a_ at position z relates the bulk value to the difference in transfer free energy of the two protonation states from bulk water to z:

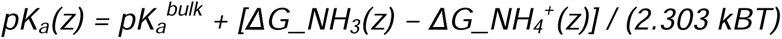

where 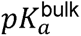 is the experimental bulk value of 9.25^43^ and both PMFs are zeroed in bulk solvent. Because no titratable proton is simulated explicitly, its free energy cancels between the two legs, and the resulting shift reflects the differential electrostatic and desolvation environment experienced by the two species along the channel.

## Results

### Cross-series comparability of independent SSME datasets

To assess cross-series comparability, matched H_2_O measurements at LPR5 were compared between Series 1 and Series 2 for WT NeRh50 and each pore variant (Figure 1). Maximum amplitude did not differ significantly between series for WT, D162A, H170A or H170D (WT: estimate −0.20 nA, 95% CI: −0.51-0.12 nA, p = 0.722; D162A: estimate −0.03 nA, 95% CI: −0.26-0.20 nA, p = 1.000; H170A: estimate −0.08 nA, 95% CI: −0.49-0.33 nA, p = 0.998; H170D: estimate −0.28 nA, 95% CI: −0.57-0.01 nA, p = 0.258; all p-values Šidák-corrected across the five comparisons). H170E, however, showed a modest but significant series effect (estimate −0.74 nA, 95% CI: −1.28-−0.20 nA, p = 0.037). This difference was not interpreted as a mechanistic effect, but as an expected batch-dependent amplitude effect for this high-current variant; consequently, series 1 and 2 datasets were not pooled.

**Figure 1.**
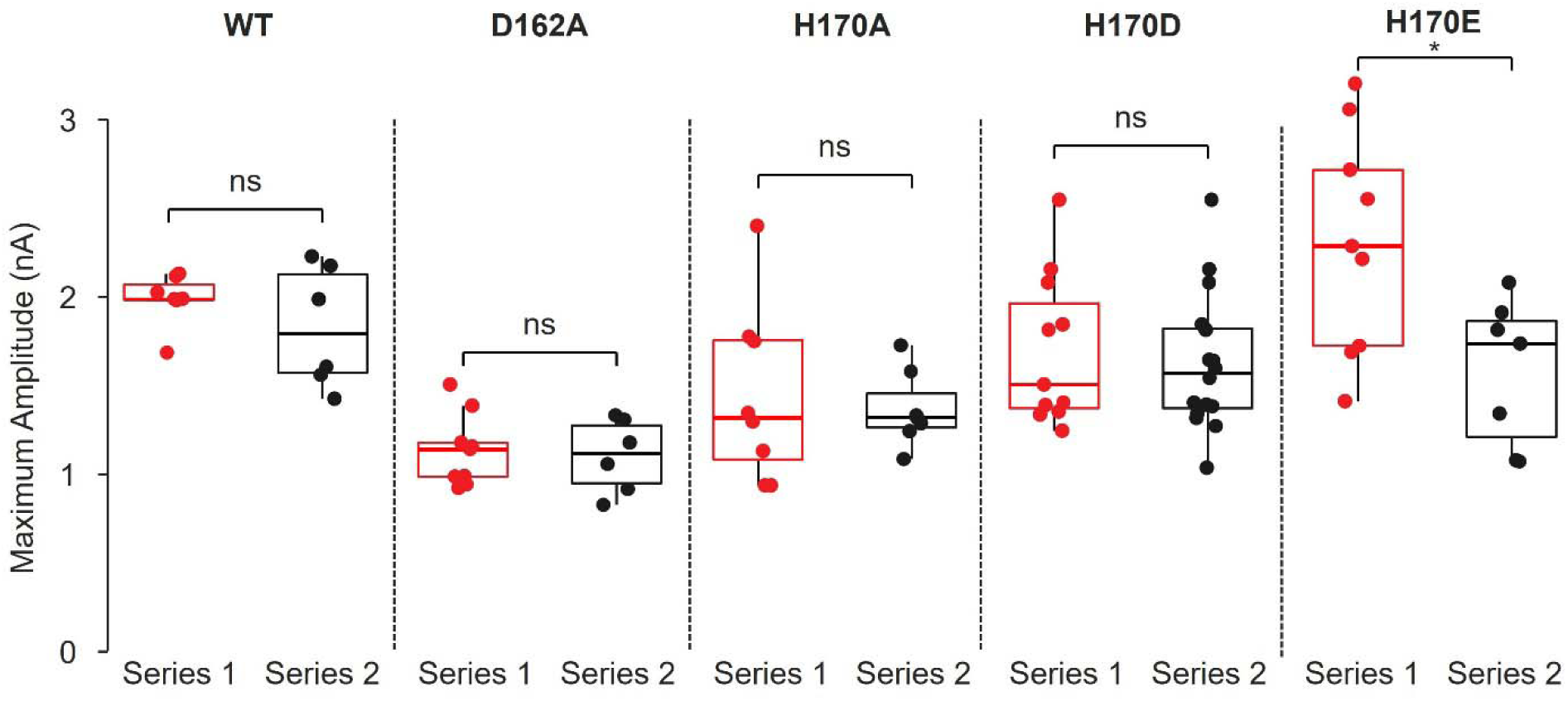
Cross-series comparison of NH_4_ ^+^-induced maximum amplitude for WT NeRh50 and pore variants. Maximum amplitude for NH_4_ ^+^-induced currents in H_2_O at LPR5, compared between Series 1 (red) and Series 2 (black), for WT NeRh50, D162A, H170A, H170D and H170E. Boxes show median and interquartile range with individual replicates overlaid; whiskers extend to the most extreme value within 1.5×IQR. Each genotype was modelled separately using a linear model with heteroskedasticity-consistent (sandwich) standard errors; pairwise comparisons between series were obtained from the model’s estimated marginal means, with Šidák correction for multiple comparisons across the five genotypes. Asterisks indicate significance: *p < 0.05, ns, not significant.

### NeRh50 retains a D_2_O-resistant electrogenic component

NH_4_^+^ deprotonation during transport has been proposed as a conserved mechanistic feature of the Amt/Mep/Rh superfamily^44^. This idea was incorporated into the two-lane model for AmtB, in which NH_4_^+^ is deprotonated and the released H^+^ moves inward through a hydrogen-bonded water network^20^. D_2_O provides a kinetic isotope test of this mechanism, because proton/deuteron transfer through hydrogen-bonded water wires is slower in D_2_O than in H_2_O. In this model, electrogenic transport depends on proton movement through a water wire, and substitution of H_2_O with D_2_O should strongly reduce or abolish the current. Consistent with this prediction, AmtB activity was previously shown to be abolished in D_2_O^20^. We therefore asked whether NeRh50 follows the same D_2_O-sensitive transport regime. Application of a 200 mM NH_4_^+^ pulse to WT NeRh50 proteoliposomes induced transient currents in H_2_O at all LPRs tested (Figure 2A). In H_2_O, maximum amplitude was reduced by 0.763 nA at LPR10 relative to LPR5 (95% CI: 0.328–1.20 nA, p = 0.0012), and by a further 0.648 nA at LPR50 relative to LPR10 (95% CI: 0.166–1.13 nA, p = 0.0087). The decay rate constant k also decreased from LPR5 to LPR10 by 23.5 s^-1^ (95% CI: 1.65–45.3 s^-1^, p = 0.034), and decreased by 20.5 s^-1^ from LPR10 to LPR50 (95% CI: −3.68–44.7 s^-1^, p = 0.106) (Figure 2A-B). This LPR-dependent behaviour is consistent with a full translocation cycle rather than isolated substrate binding or local charge displacement^45^. We next tested whether this transport-associated current was abolished by D_2_O. Solvent exchange reduced the NH_4_^+^-induced response but did not eliminate it (Figure 2C-D). At LPR5, maximum amplitude decreased by 0.832 nA (95% CI: 0.454-1.21 nA, p = 0.00010) in D_2_O, leaving a residual response equivalent to ∼55% of the initial H_2_O signal. Return to H_2_O restored the response, confirming that D_2_O exposure did not irreversibly unfold the protein, damage proteoliposome integrity or sensor capacitance (Figure 2C-D). In contrast to maximum amplitude, the decay rate constant k at LPR5 was not significantly affected by D_2_O substitution (D_2_O vs H_2_O: 95% CI: −15.95-20.21 s^-1^, p = 0.985; D_2_O>H_2_O vs H_2_O: 95% CI: −31.33-10.63 s^-1^, p = 0.493; D_2_O>H_2_O vs D_2_O: 95% CI: −32.85-7.89 s^-1^, p = 0.317; all ns; Figure 2D), indicating that solvent exchange selectively attenuates the size of the electrogenic response without altering its kinetics. Residual NH_4_^+^-induced currents detected in D_2_O were LPR-dependent: maximum amplitude decreased by 0.232 nA (95% CI: 0.006-0.458 nA, p = 0.043) from LPR5 to LPR10, and by a further 0.481 nA (95% CI: 0.218-0.743 nA, p = 0.00063) from LPR10 to LPR50. Corresponding decay rate constants k were also LPR-dependent, decreasing by 33.6 s^-1^ (95% CI: 20.8-46.4 s^-1^, p = 1.5 × 10^-5^) from LPR5 to LPR10, and by a further 17.7 s^-1^ (95% CI: 2.9-32.5 s^-1^, p = 0.017) from LPR10 to LPR50 (Figure 2E-F). Thus, the residual D_2_O signal retains significant LPR-dependent behaviour, confirming a full translocation rather than a nonspecific artefact.

**Figure 2.**
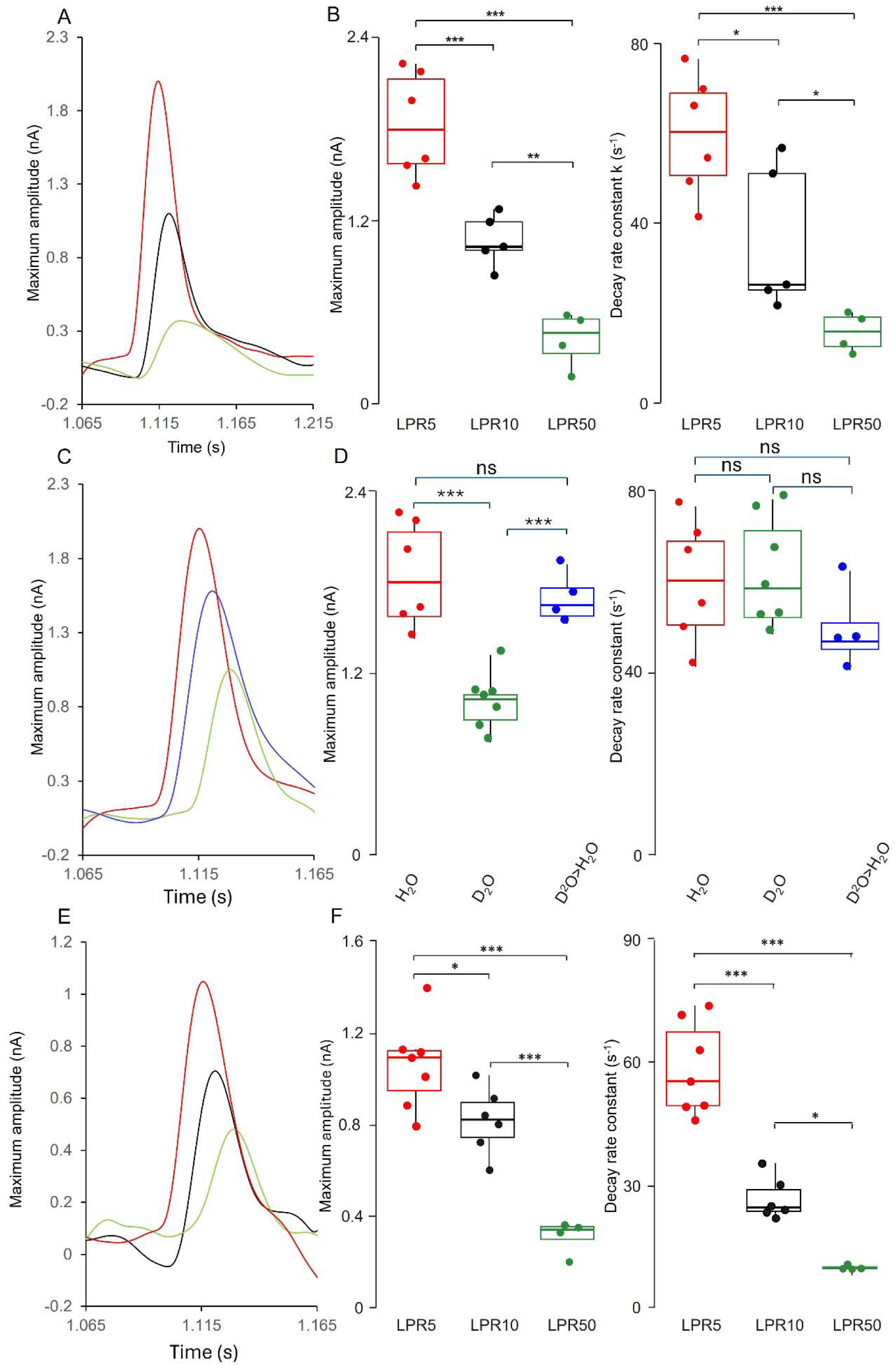
NeRh50 generates D_2_O-resistant, LPR-dependent electrogenic currents in response to NH ^+^ stimulation. **(A)** Representative SSME transient currents recorded in H_2_O after application of 200 mM NH_4_Cl to WT NeRh50 proteoliposomes reconstituted at LPR5 (red), LPR10 (black) and LPR50 (green). **(B)** Quantification of H_2_O measurements showing maximum amplitude and decay rate constant k across LPR5, LPR10 and LPR50. Boxes show median and interquartile range with individual replicates overlaid; whiskers extend to the most extreme value within 1.5×IQR. LPR was modelled as the sole predictor of each response in a linear model (ordinary least squares); pairwise comparisons between LPR levels were obtained from the model’s estimated marginal means with Šidák correction for multiple comparisons. **(C)** Representative solvent-exchange experiment at LPR5 showing the initial NH_4_ ^+^-induced response in H_2_O (red), the residual response in D_2_O (green), and the response after return to H_2_O following D_2_O exposure (blue). **(D)** Quantification of solvent-exchange measurements at LPR5 showing maximum amplitude and decay rate constant k in H_2_O, D_2_O and after return to H_2_O (D_2_O>H_2_O). Condition was modelled as the sole predictor in a linear model; pairwise comparisons between conditions used the model’s estimated marginal means with Šidák correction. **(E)** Representative NH_4_ ^+^-induced responses recorded in D_2_O at LPR5 (red), LPR10 (black) and LPR50 (green). **(F)** Quantification of D_2_O measurements showing maximum amplitude and decay rate constant k across LPR5, LPR10 and LPR50, modelled and tested as in (B). Asterisks indicate significance: *p < 0.05, **p < 0.01, ***p < 0.001; ns, not significant.

### D162 couples NH_4_^+^ engagement to inward uptake but is dispensable for substrate export

Following a 200 mM ammonium pulse, NeRh50 D162A produced a measurable transient current. At LPR5 this was smaller than that of WT by 0.892 nA (95% CI: 0.674-1.110 nA, p = 2.95 × 10^-12^; Figure 3A-C). The D162A response showed an estimated decrease of 0.183 nA as protein density was lowered from LPR5 to LPR10, although this did not reach significance (95% CI: −0.030-0.395 nA, p = 0.125; Figure 3A-C), indicating that NH_4_^+^ still engages the transporter in a protein-dependent manner. However, these amplitude changes were much smaller than in WT, indicating that D162A weakens the coupling between substrate engagement and productive transport. The clearest defect was observed in the decay kinetics (Figure 3A-B, D). In WT NeRh50, k decreased by 11.0 s^-1^ (95% CI: 0.415-21.63 s^-1^, p = 0.038) as protein density fell from LPR5 to LPR10, consistent with a full translocation cycle. In D162A, k did not decrease significantly between LPR5 and LPR10 (+1.86 s^-1^, 95% CI −9.39 to 13.11 s^-1^, p = 0.998). Thus, although NH_4_^+^ still produces a protein-dependent signal in D162A, the signal no longer shows the WT-like kinetic signature of efficient translocation. Michaelis-Menten analysis supported this conclusion: D162A retained a measurable apparent K_m_ for NH_4_^+^, but this increased to 250.60 ± 62.80 mM, compared with 59.30 ± 8.50 mM for WT NeRh50 (Figure 3E). Solvent exchange further showed that the residual D162A signal was insensitive to D_2_O (Figure 3F). At LPR5, maximum amplitude was 1.10 ± 0.21 nA in H_2_O, 0.81 ± 0.38 nA in D_2_O and 1.18 ± 0.35 nA after return to H_2_O. The H_2_O-to-D_2_O decrease was not significant and recovery was complete, confirming that the solvent exchange was reversible (Figure 3F). Together, these data indicate that D162A retains a residual NH_4_^+^-associated electrogenic response but uncouples pore-entry engagement from inward translocation.

We next tested whether the residual SSME signal of NeRh50 D162A corresponded to inward ammonium uptake *in vivo*. In the triple-mepΔ yeast strain, both WT AmtB and WT NeRh50 restored growth at 0.5 and 1 mM NH_4_^+^, whereas AmtB D160A and NeRh50 D162A, failed to complement (Figure 4A). This parallel loss-of-function phenotype is consistent with previous results obtained for AmtB D160A under different basal medium conditions^20^. At 3 mM NH_4_^+^, all strains grew after 4 days, including the pDR195 empty-vector control, indicating that growth at this concentration reflects NH_3_ diffusion and/or uptake through endogenous nonspecific transporters and/or channels. Because the complementation assay was performed at 0.5-1 mM NH_4_^+^, far below the apparent K_m_ values measured for both WT NeRh50 and NeRh50 D162A, the failure of D162A to complement cannot be explained by reduced apparent affinity alone. Instead, it confirmed a specific defect in coupling NH_4_^+^ engagement to inward translocation. Methylammonium (MeA) is a toxic ammonium analogue in yeast^20,30,31^, and Rh proteins expressed in yeast have been associated with MeA-resistant phenotypes consistent with substrate export/protection^46^. At 30 and 100 mM MeA, pDR195 still grew, indicating that these conditions were not sufficiently selective to define NeRh50-specific protection. The clearest discriminatory condition was 200 mM MeA. Under this condition, pDR195, WT AmtB and AmtB D160A showed no growth, whereas WT NeRh50 retained growth (Figure 4A). This indicates that MeA resistance is not a generic consequence of expressing an ammonium transporter but is associated with NeRh50. Importantly, NeRh50 D162A also retained growth at 200 mM MeA despite failing to complement growth on limiting NH_4_^+^. We next asked whether the NeRh50-dependent MeA-protective phenotype persisted in WT yeast, where endogenous Mep transporters provide a controllable route for MeA entry (Figure 4B). Proline and glutamate were used as Mep-permissive nitrogen sources, with proline providing the stronger Mep-inducing condition, whereas glutamine represses Mep expression. Glutamine plus ammonium was included as a restrictive control, combining Mep repression with ammonium competition for MeA transport. The pDR195 control defined the MeA-entry conditions in this assay. At 30 mM MeA, pDR195 grew on glutamine but was inhibited on proline and glutamate, consistent with Mep-dependent MeA toxicity. At higher MeA concentrations, pDR195 also became sensitive on glutamine, consistent with background MeA entry, whereas ammonium partially rescued growth under glutamine plus ammonium. WT NeRh50 protected cells under conditions where pDR195 was sensitive, including proline and glutamate at 30 mM MeA and glutamine at higher MeA concentrations. This protection was not observed with WT AmtB or AmtB D160A, showing that MeA protection is specific to NeRh50 and is not a generic consequence of ammonium-transporter expression. NeRh50 D162A retained a clear MeA-protective phenotype despite its loss of inward NH_4_^+^ complementation. Protection was evident on glutamate at 30 mM MeA and on glutamine at 100-200 mM MeA, but not under the stronger proline-induced MeA-entry condition (Figure 4B). Thus, D162A preserves an MeA-protective/export-associated output that can be detected when MeA influx is not maximal. Together with the triple-mepΔ assay, these data show that D162 is required for inward NH_4_^+^ uptake but is dispensable for the NeRh50-associated MeA-protective/export activity, demonstrating that inward uptake and MeA-protective/export-associated handling can be decoupled in NeRh50.

**Figure 3.**
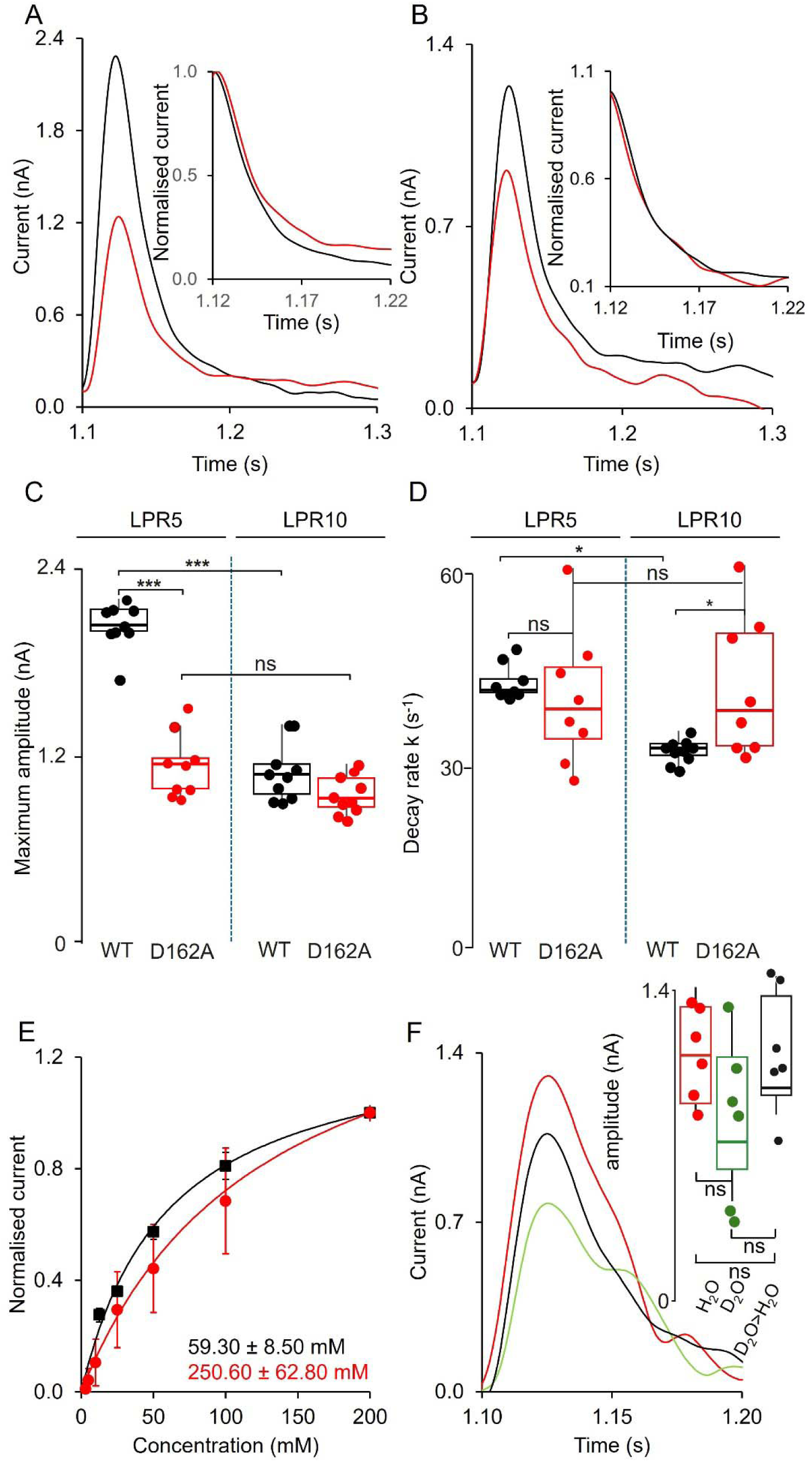
D162A uncouples ammonium binding from translocation. **(A, B)** Representative SSME current traces for WT (black) and D162A (red) NeRh50 upon an NH_4_^+^ 200 mM pulse, recorded at LPR5 (A) and LPR10 (B). **Insets** show peak-normalised traces over the decay phase. **(C)** Maximum current amplitude and **(D)** decay rate constant for WT and D162A at LPR5 and LPR10. Individual data points are overlaid on box plots showing the median and interquartile range, with individual replicates overlaid; whiskers indicate the most extreme data points within 1.5× the interquartile range; WT in black, D162A in red. Statistical comparisons by linear model with Šidák-corrected pairwise contrasts (emmeans); ***p < 0.001, **p < 0.01, *p < 0.05, ns = not significant. **(E)** NH_4_^+^ concentration-response curves for WT (black squares) and D162A (red circles) normalised current, fitted with the Michaelis-Menten equation. Error bars, SEM. **(F)** Effect of D_2_O substitution on D162A current. Traces recorded in H_2_O (red), D_2_O (green), and after D_2_O-to-H_2_O exchange (D_2_O>H_2_O, black). Boxes show the median and interquartile range, with individual replicates overlaid; whiskers indicate the most extreme data points within 1.5× the interquartile range. **Inset:** maximum current amplitude under each condition; pairwise comparisons by linear model with Šidák-corrected pairwise contrasts, not significant (ns).

**Figure 4.**
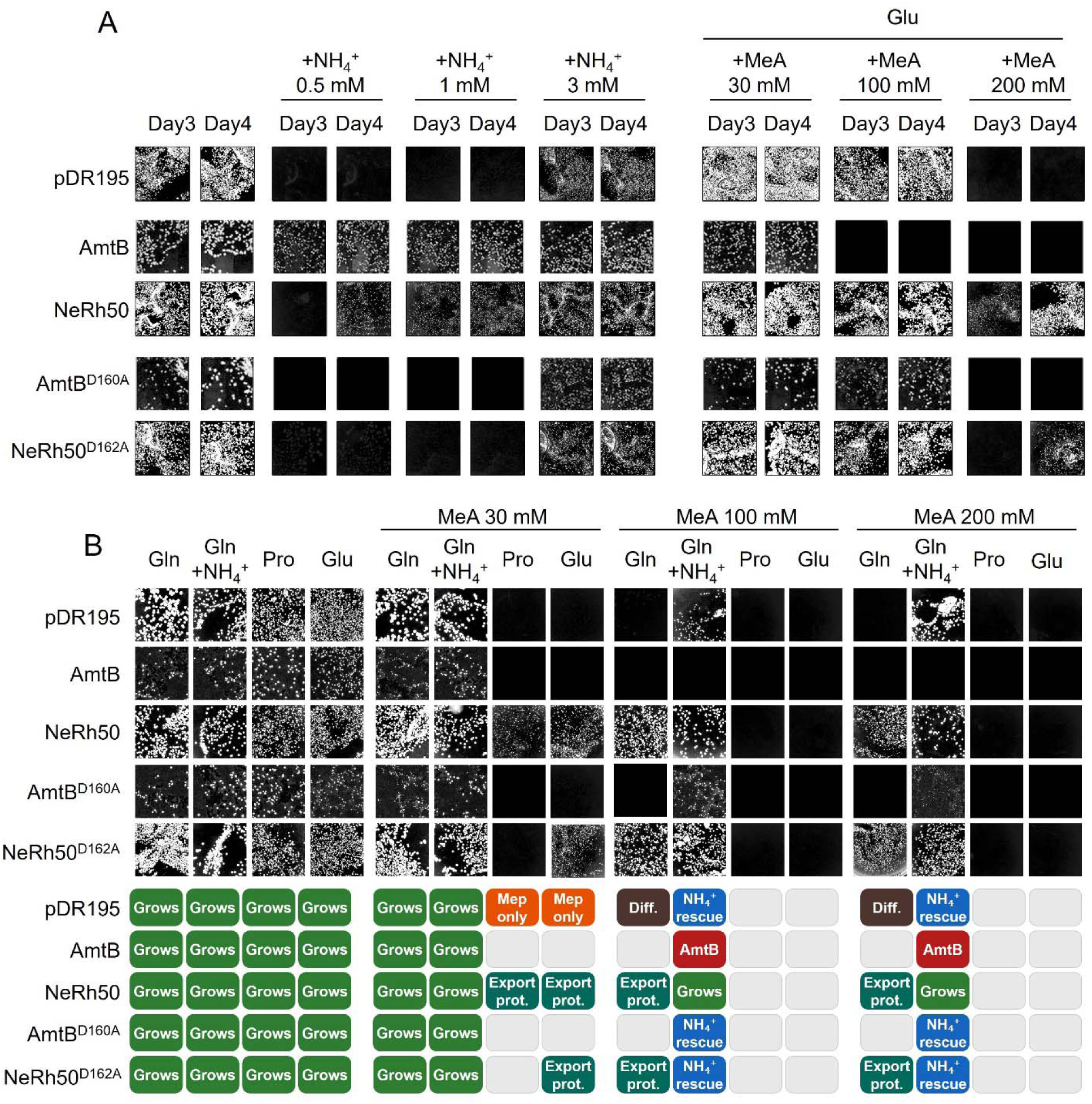
Yeast assays show that NeRh50 D162A loses inward NH_4_^+^ uptake but retains MeA-protective activity. **(A)** Triple-mepΔ yeast complementation and MeA-resistance assay. S. cerevisiae triple-mepΔ cells expressing pDR195 empty vector, WT AmtB, WT NeRh50, AmtB D160A or NeRh50 D162A were grown on glutamate 0.2% medium as a growth control, on limiting NH_4_^+^ medium to test inward ammonium uptake, or on glutamate medium supplemented with methylammonium (MeA) to test MeA resistance. Plates were imaged after 3 and 4 days of growth. **(B)** MeA toxicity assay in a WT yeast background, where endogenous Mep-dependent MeA entry is controlled by nitrogen source. Cells expressing the indicated transporters were grown on glutamine 0.1% (Gln), glutamine plus 20 mM ammonium (Gln+NH_4_^+^), proline 0.1% (Pro) or glutamate 0.1% (Glu), in the absence or presence of MeA. The lower panel provides a categorical interpretation of the growth phenotypes, assigned by comparison with the relevant control under the same condition. Green squares indicate growth; grey squares indicate no detectable or non-informative growth; orange squares indicate MeA toxicity dependent on endogenous Mep activity; brown squares indicate toxicity consistent with MeA diffusion/background entry; blue squares indicate NH_4_ ^+^ rescue; red squares indicate transporter-associated MeA sensitivity; and teal squares indicate MeA-protective/export.

### H170 is essential for export but not import activity

We next examined H170, the first histidine of the conserved twin-His motif. This position corresponds to AmtB H168 and is central to proposed proton-transfer models in which histidine chemistry, tautomeric state and water-wire organisation contribute to proton relay^20,23^. Application of 200 mM NH_4_^+^ to NeRh50 H170A, H170D and H170E proteoliposomes produced clear NH_4_^+^-induced transient LPR-dependent responses (Figure 5A-B). At LPR5, WT NeRh50 showed a maximum amplitude of 2.03 ± 0.15 nA, compared with 1.45 ± 0.50 nA for H170A, 1.70 ± 0.42 nA for H170D and 2.36 ± 0.63 nA for H170E. Thus, H170 substitution did not abolish NH_4_^+^-associated electrogenic activity (Figure 5A-B). At LPR5, all H170 variants had reduced decay rate constants in comparison to WT NeRh50: H170A −8.451 s^-1^ (95% CI −14.05 to −2.85, p = 0.0015); H170D −12.699 s^-1^ (95% CI −17.52 to −7.88, p = 9.38 × 10^-8^); H170E −20.319 s^-1^ (95% CI −25.56 to −15.08, p = 1.42 × 10^-12^) (Figure 5B). These data show that these H170 variants translocate NH_4_^+^. Solvent exchange exposed a sharp loss of the D_2_O-resistant component across all H170 variants (Figure 5C). H170A retained only a small fraction of its H_2_O amplitude in D_2_O (1.37 ± 0.22 nA to 0.39 ± 0.05 nA), a loss that fully reversed on return to H_2_O (1.66 ± 0.33 nA). For H170E, no discernible current was observed in D_2_O; recovery in H_2_O was complete (1.96 ± 0.22 nA, Figure 5C). For H170D, a small residual D_2_O response was measurable (0.36 ± 0.05 nA, significantly lower than the H_2_O response; 95% CI 0.723-1.396 nA, p = 1.79 × 10^-7^), and recovery in H_2_O was partial and numerically lower than the initial H_2_O amplitude, although this difference did not reach significance (1.22 ± 0.27 nA versus 1.42 ± 0.19 nA; 95% CI −0.057-0.453 nA, p = 0.160; Figure 5C). H170 is therefore not required for transport itself but is essential for the D_2_O-resistant pathway that distinguishes NeRh50 from canonical AmtB. Michaelis-Menten analysis further showed that H170 substitutions did not abolish substrate access to the pore (Figure 5D). The apparent K_m_ values were 303.20 ± 25.20 mM for H170A, 144.50 ± 37.70 mM for H170D and 64.70 ± 21.70 mM for H170E, compared with 59.30 ± 8.50 mM for WT NeRh50 (Figure 5D). Thus, all three variants retain measurable NH_4_^+^ concentration-dependent responses. H170A and H170D showed increased apparent K_m_ values. H170E retained a WT-like apparent K_m_ and showed an increased maximum amplitude, indicating that this substitution increases ammonium flux, like the equivalent mutant in H168E in AmtB^20^.

Remarkably, a D_2_O-resistant signal persists in D162A but is lost in the H170 variants, even though D162 remains intact. This difference is explained by where each mutation acts relative to the downstream transport cycle, including the twin-His region implicated in water-wire-mediated proton transfer. In D162A, substrate engagement is interrupted at the pore-entry step: NH_4_ ^+^ can still bind or engage the external vestibule, but this event is poorly coupled to progression into the downstream pore. The residual signal therefore reports an upstream, entry-associated charge displacement rather than solvent-coupled translocation, explaining why it is not strongly affected by D_2_O. In the H170 variants, by contrast, D162-dependent pore-entry engagement is preserved and NH_4_ ^+^-associated transport proceeds into the downstream cycle. However, substitution of H170 removes the Rh-specific pore state that supports the D_2_O-resistant component in WT NeRh50. The remaining transport output is therefore restricted to a D_2_O-sensitive, proton-transfer-dependent component, similar to the canonical AmtB-like pathway. Thus, loss of the D_2_O-resistant signal in H170A, H170D and H170E does not indicate failure of substrate recruitment at D162; it shows that D162-dependent entry must be coupled to an intact H170-controlled pore state to generate the proton-transfer-independent component of NeRh50 transport.

To test whether the H170-dependent SSME phenotype was reflected in cellular substrate handling, we compared WT and twin-His variants in yeast complementation and MeA-resistance assays (Figures 6 and 7). As shown above (Figure 4), pDR195 did not complement growth on limiting NH_4_^+^ whereas WT AmtB and WT NeRh50 restored growth, with WT AmtB remaining MeA-sensitive and WT NeRh50 retaining growth at high MeA. NeRh50 H170A, H170D and H170E all restored growth on NH_4_^+^ relative to pDR195, particularly by day 4 and at higher NH_4_^+^ concentrations. By contrast, all three variants showed impaired growth at elevated MeA: H170A showed only weak or delayed growth at 30 mM MeA and failed at higher concentrations, while H170D and H170E were strongly impaired across the entire MeA series. The equivalent AmtB variants showed a different pattern. AmtB H168A retained ammonium complementation and remained MeA-sensitive, similar to WT AmtB, indicating that this substitution preserves sufficient inward ammonium transport without conferring NeRh50-like MeA protection. By contrast, acidic substitutions at AmtB H168 caused a strong growth defect under ammonium conditions, particularly at 3 mM NH_4_^+^, where the pDR195 control was able to grow. This phenotype is consistent with previous work showing that acidic substitution of AmtB H168 increases ammonium-associated flux and can convert regulated transporter activity into a dysregulated, channel-like conductance, thereby causing ammonium-dependent toxicity rather than productive growth complementation^20,31^. Thus, the growth defect of AmtB H168D/E is best interpreted as toxic over-entry or loss of transport control, not as absence of ammonium movement.

In the WT yeast background, all strains grew on the permissive nitrogen sources in the absence of MeA, confirming that expression of AmtB, NeRh50 and the H168/H170 variants was not generally toxic (Figure 7). As expected, MeA inhibited growth under conditions where endogenous Mep-dependent uptake contributes to toxicity, whereas addition of NH_4_^+^ partially rescued growth of the empty-vector control. WT AmtB did not protect cells from MeA toxicity, consistent with MeA entry through AmtB, whereas WT NeRh50 supported growth under MeA stress. NeRh50 H170A, H170D and H170E all grew normally in the absence of MeA, but none retained the MeA-resistance phenotype of WT NeRh50; growth under MeA was weak, condition-dependent, or dependent on NH_4_^+^ rescue. Consistent with the complementation assay above, AmtB H168A, H168D and H168E also failed to confer MeA resistance and remained MeA-sensitive like WT AmtB. In addition, the acidic H168D/E substitutions showed NH_4_^+^-dependent growth defects, consistent with previous reports that these mutations increase ammonium flux and can cause ammonium toxicity rather than productive complementation^20,31^. Thus, AmtB H168D/E behave as toxic high-flux import variants, whereas NeRh50 H170 variants retain import but lose the Rh-like MeA-protective/export activity. Together, these data show that in NeRh50, H170 is dispensable for basal growth in the absence of MeA but is required for the Rh-like MeA-resistant/export activity, corroborating the SSME-based dissociation between import and the D_2_O-resistant/export-activity.

**Figure 5.**
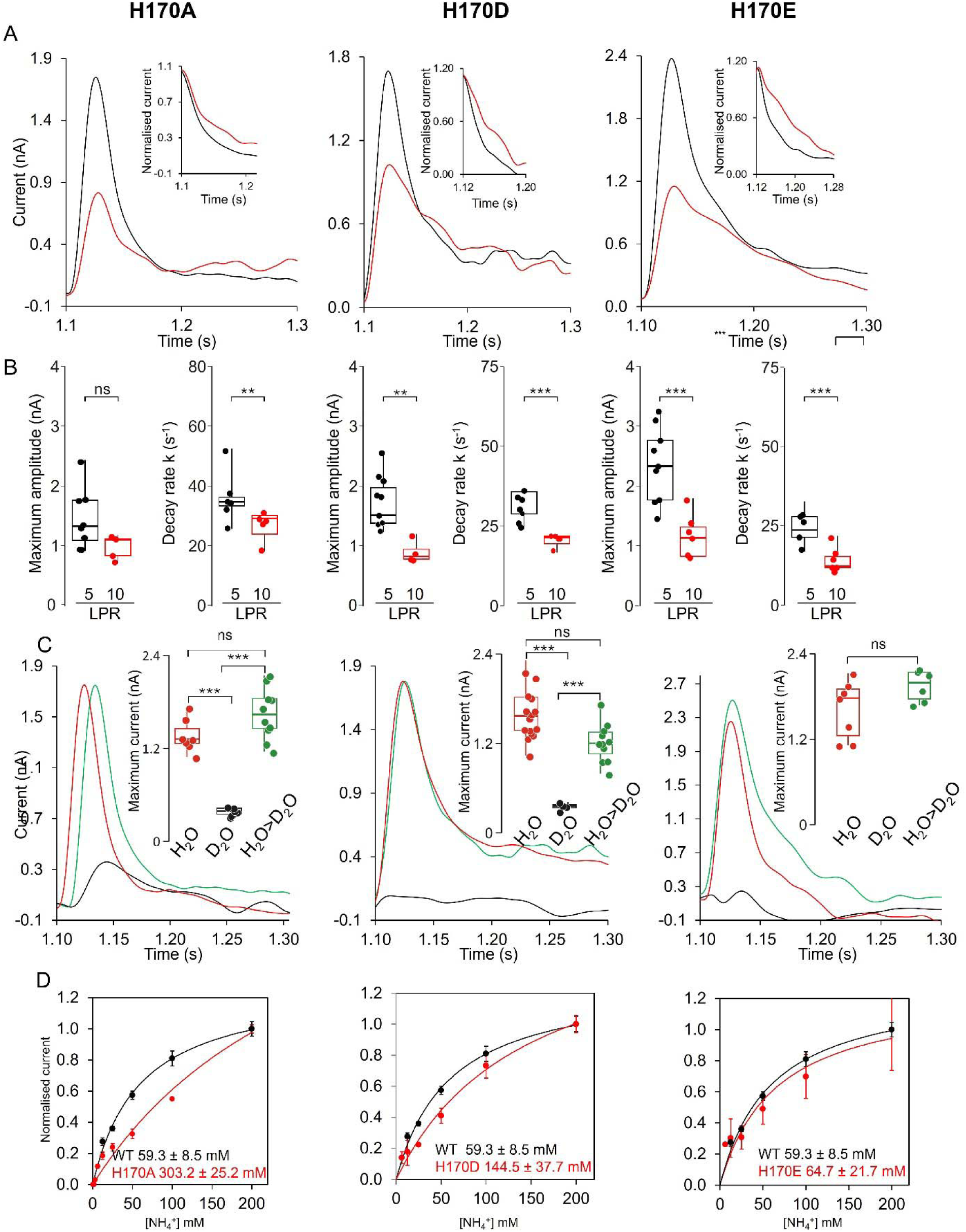
Electrophysiological characterisation of H170 variants. **(A)** Representative SSME transient currents recorded from NeRh50 H170A, H170D and H170E proteoliposomes following application of 200 mM NH_4_Cl at pH 7.0. Currents were recorded at LPR5 (black) and LPR10 (red). Insets show normalised current decays used to compare kinetic behaviour. **(B)** Quantification of NH_4_ ^+^-induced maximum amplitude and decay rate constant k for each H170 variant at LPR5 (black) and LPR10 (red). Individual data points are overlaid and box plots show the median and interquartile range, with individual replicates overlaid; whiskers indicate the most extreme data points within 1.5× the interquartile range. Statistical comparisons by linear model with Šidák-corrected pairwise contrasts (emmeans); ***p < 0.001, **p < 0.01, *p < 0.05, ns = not significant. **(C)** Effect of D_2_O substitution on H170 variant current. Traces recorded in H_2_O (red), D_2_O (black), and after D_2_O-to-H_2_O exchange (D_2_O>H_2_O, green). Inset: maximum current under each condition, presented as in (B). Statistical comparisons by linear model with Šidák-corrected pairwise contrasts. For H170E, no quantifiable D₂O peak was detected; the D₂O trace is shown, but the condition was excluded from the inset and statistical comparison. **(D)** NH_4_ ^+^ concentration-response curves for WT (black) and each H170 variant (red), normalised current, fitted with the Michaelis-Menten equation. Apparent K_m_ ± SEM is indicated on each panel. Error bars, SEM.

**Figure 6.**
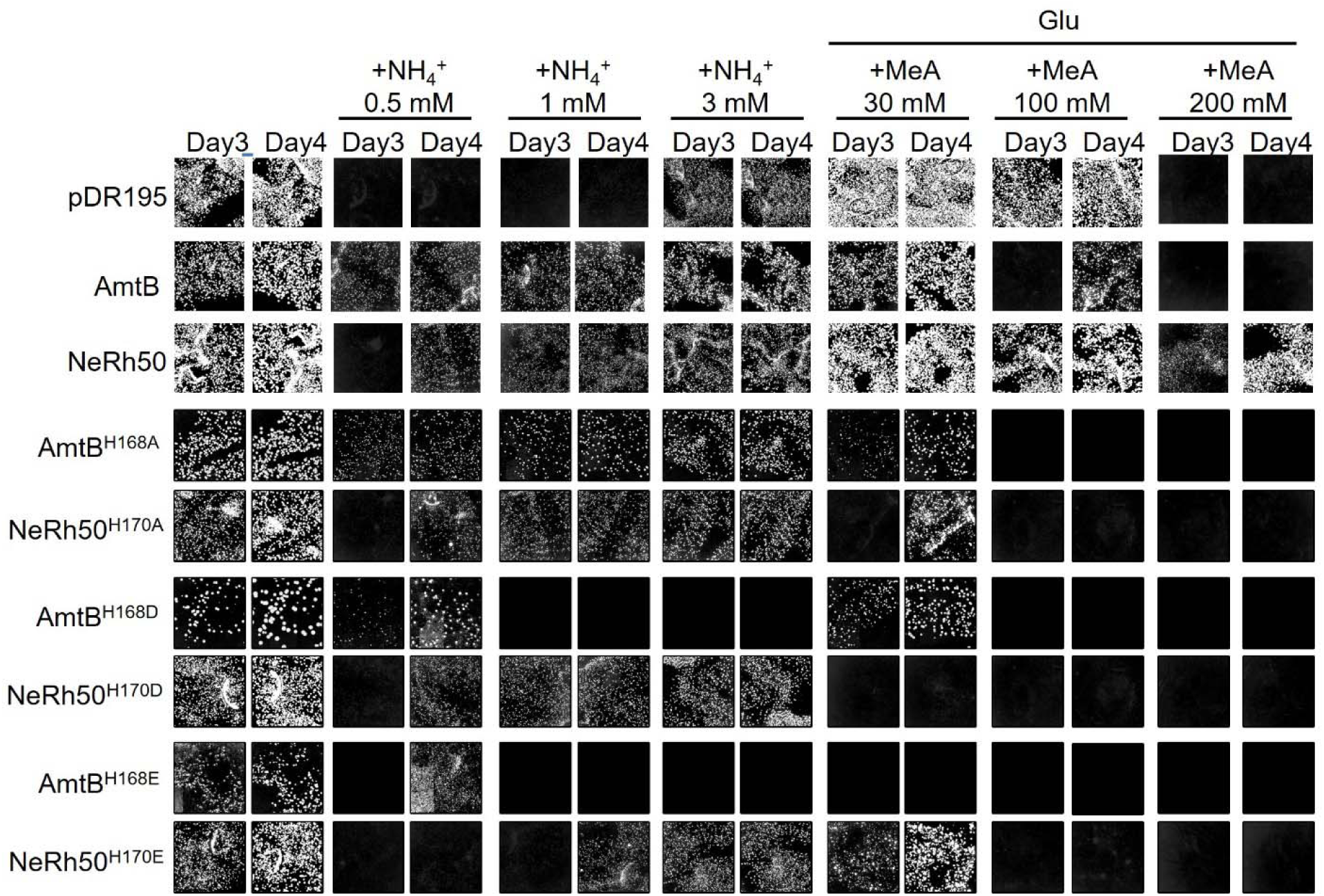
H170 variants retain import but lose export activity. Growth assay of S. cerevisiae triple-mepΔ cells expressing pDR195 empty vector, WT AmtB, WT NeRh50, AmtB H168A/D/E or the equivalent NeRh50 H170A/D/E variants were grown on glutamate 0.2% medium as a growth control, on limiting NH_4_ ^+^ medium to test inward ammonium uptake, or on glutamate medium supplemented with methylammonium (MeA).

**Figure 7.**
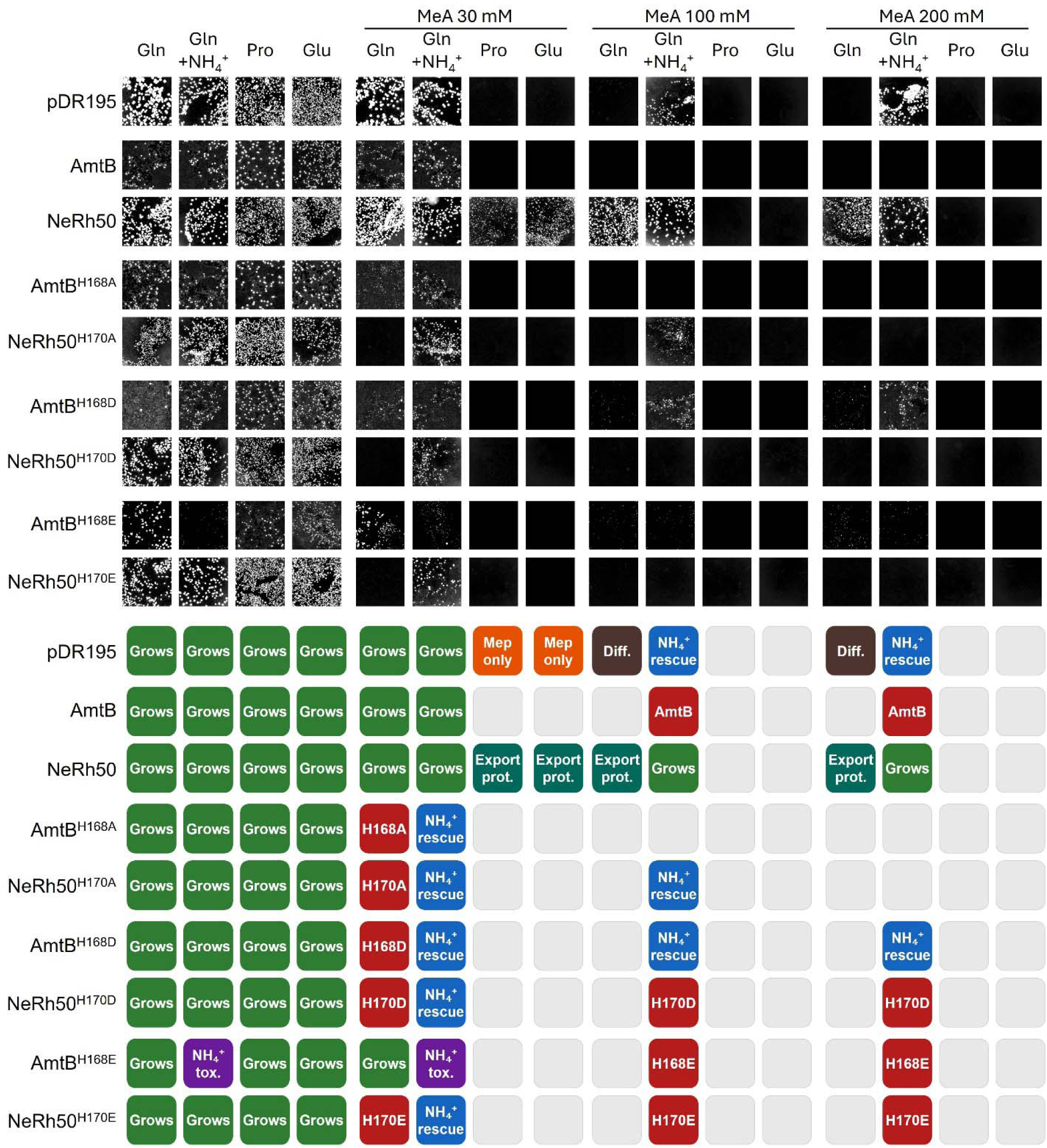
MeA resistance in WT yeast expressing NeRh50 H170 and AmtB H168 variants. MeA toxicity assay in a WT yeast background, where endogenous Mep-dependent MeA entry is controlled by nitrogen source. Cells expressing the indicated transporters were grown on glutamine 0.1% (Gln), glutamine plus 20 mM ammonium (Gln+NH_4_^+^), proline 0.1% (Pro) or glutamate 0.1% (Glu), in the absence or presence of MeA. Lower panel: Heatmap summary of MeA phenotypes across all strain-condition combinations. Colour coding reflects the mechanistic interpretation of each phenotype: green (Grows), normal growth; teal (Export prot.), MeA-protective growth requiring active heterologous export under conditions where Mep-mediated MeA entry is active (Pro, Glu) or diffusion-limited (Gln); orange (Mep only), MeA toxicity attributable to endogenous Mep activity; blue (NH_4_ ^+^ rescue), growth restored by ammonium competition for transport/metabolism under Gln+NH_4_ ^+^; purple (NH_4_ ^+^ tox.), growth inhibition by ammonium itself with hyperactive inward transport AmtB H168E^31^; dark red (Diff.), toxicity due to MeA diffusion; red, variant-specific intermediate phenotype; grey, no growth.

**Figure 8.**
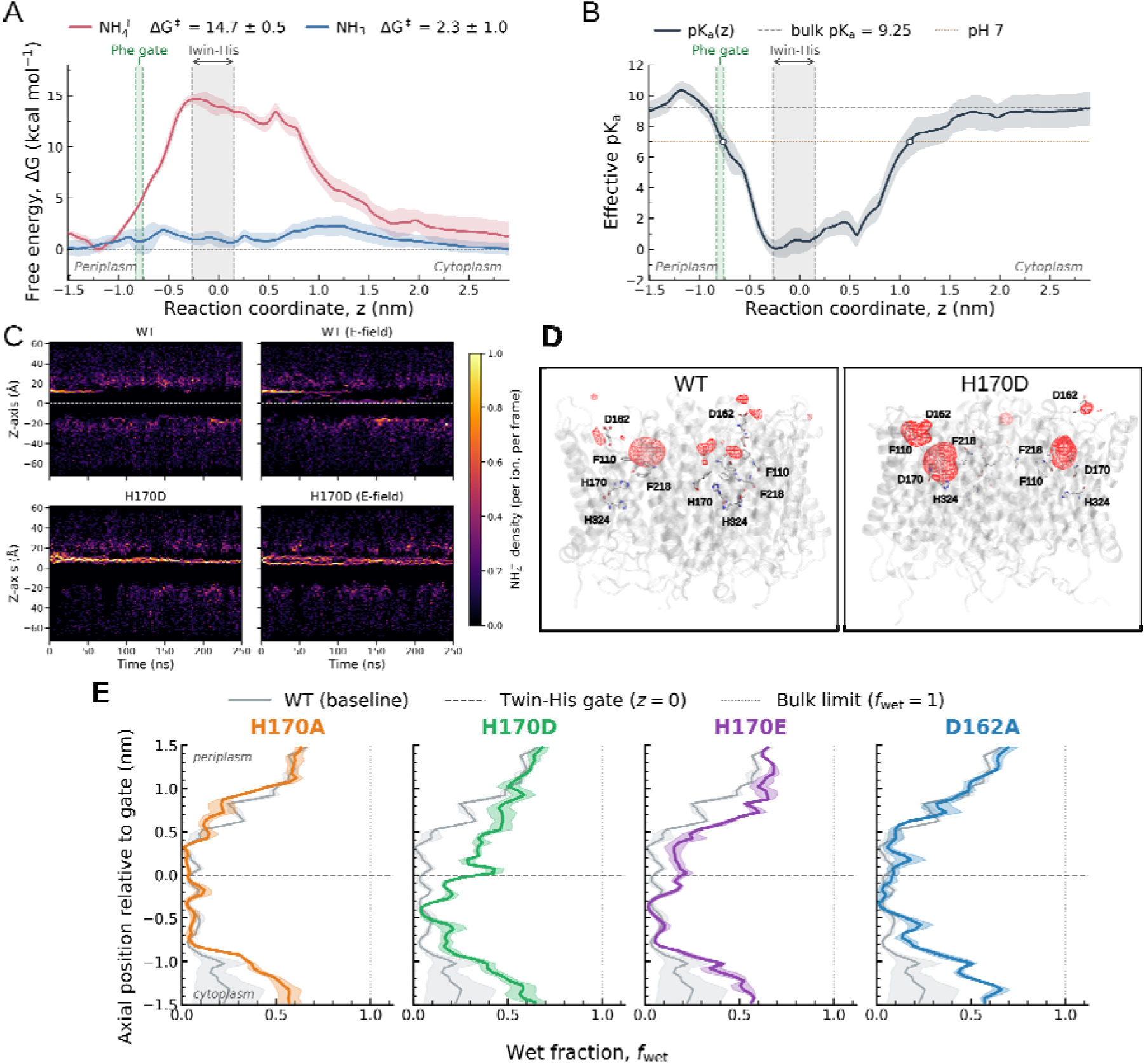
Molecular simulations link WT pore dehydration to NH ^+^ destabilisation and H170D-dependent trapping. **(A)** Axial potentials of mean force for fixed-state NH ^+^ and NH through one WT NeRh50 protomer. Solid lines show the mean profiles; shading shows the reported uncertainty and ΔG‡ values give the maximum barriers. The Phe-gate and twin-His constriction are indicated. **(B)** Position-dependent effective substrate pK_a_ derived from the two free-energy profiles. Horizontal lines mark the aqueous bulk pK_a_ of 9.25 and pH 7. **(C)** Time-resolved axial NH ^+^ density in WT and H170D, with and without a 0.05 V nm^-1^ field applied along −z. Densities were pooled across three 250-ns replicates and normalised per ion and frame; the dashed line marks z = 0. Late cytoplasmic-face density in WT reflects periodic-boundary crossing through bulk solvent rather than pore translocation. **(D)** Time-averaged three-dimensional NH ^+^ density in WT and H170D. Red meshes show regions of occupancy at a common isovalue. **(E)** Axial wet-fraction profiles from three 500-ns substrate-free replicates of H170A, H170D, H170E and D162A relative to WT. Lines show the mean profiles and shading shows the associated variation. For panels A and B, negative z values are periplasmic; for panels C and E, positive z values are periplasmic. Residues are labelled using experimental numbering.

### Molecular simulations link WT pore dehydration to NH_4_^+^ destabilisation and H170D-dependent trapping

To determine whether WT NeRh50 can accommodate ammonium (NH_4_^+^), ammonia (NH_3_), or both, we used umbrella sampling to compare their axial potentials of mean force through the central pore of one WT NeRh50 protomer. The profiles differed markedly: NH_4_^+^ encountered a maximum barrier of 14.7 ± 0.5 kcal mol^-1^, whereas the corresponding barrier for NH_3_ was only 2.3 ± 1.0 kcal mol^-1^ (Figure 8A). Focusing on the NH_4_^+^ profile in more detail, moving from the periplasmic side to the cytoplasmic side, the first notable feature is a shallow NH_4_^+^ minimum near z ≈ −1.2 nm corresponding to residue D162, which has previously been hypothesized as a periplasmic recruitment region. This interpretation is consistent with work implicating the AmtB periplasmic vestibule in NH_4_^+^ recruitment^47^ and identifying the homologous D160 residue as a key determinant of recruitment^48^. The second feature of interest is the aforementioned energetic barrier at *z* ≈ −0.3 nm, within the twin-His constriction formed by H170 and H324. To further investigate these regions, we then performed pK_a_ calculations and conventional molecular dynamics (MD) simulations, both at equilibrium and with an external constant electric field. Within the periplasmic vestibule (*z* ≈ −1.5 to −1.0 nm), the effective pK_a_ increased from the aqueous bulk value of 9.25^43^ to ∼10.8 at the recruitment minimum. It then fell sharply across the Phe-gate and reached a minimum of ∼0.3 in the central, twin-His region (Figure 8B). These data suggest that WT NeRh50 may recruit NH_4_^+^ at the periplasmic entry but disfavors a persistently charged substrate in the central pathway. Next, we explored the dynamics of NH_4_^+^ entry into WT NeRh50 and the single point mutant H170D. We performed two types of MD simulations in which substrates were placed in the bulk solvent, equilibrium MD and with a constant electric field applied across the simulation box. Three NH_4_^+^ ions were initially placed in the periplasm-facing vestibule of the WT and mutant proteins (one ion per protomer). Two-dimensional NH_4_^+^ density maps as a function of time, calculated from both equilibrium and electric field simulations show initial occupancy of NH_4_^+^ near D162 in WT at t = 0–60 ns, after which this region is no longer occupied (Figure 8C). Neither the equilibrium nor field-driven 250-ns trajectories showed persistent NH_4_^+^ occupancy in the central pore. These results indicate that WT NeRh50 strongly disfavors stable NH_4_^+^ occupancy in the central pore on the simulated timescale. However, they do not exclude transient passage of NH_4_^+^. In contrast, H170D produced a strong density population around the introduced Asp170 side chain. The corresponding occupancy persisted for almost the entire 250-ns trajectory, with a maximum dwell time of 244 ns under the applied field (Supplementary Figure S1). Three-dimensional plots (Figure 8D) of average NH_4_^+^ occupancy for WT and the H170D showed clear differences with NH_4_^+^ occupancy shifting towards the introduced Asp170 side chain in H170D.

Finally, we performed a set of simulations to explore the hydration properties of the WT NeRh50, H170D and three other mutants, H170A, H170E, D162A. Three independent simulations were performed for each protein in which no substrate was present. WT NeRh50 remained poorly hydrated through the central pore; similarly, H170A and D162A retained comparably dry central profiles (Figure 8E). In contrast, the wet fraction around the His-His region was increased in H170D and H170E. Pore-radius analysis showed that D162A retained WT-like central radii, whereas the acidic H170 substitutions produced a local, position-dependent enlargement extending primarily from *z* ≈ 0 to 1.0 nm on the periplasmic side of the twin-His gate (Supplementary Figure S2A). Inter-residue distance analyses showed wider His-His and Phe-Phe constrictions in H170D and H170E compared to WT. The detailed radius and distance distributions are in the Supplementary Information (Supplementary Figure S2).

Together, these simulations identify D162 as the periplasmic recruitment region and H170 as a key controller of the downstream pore state. Acidic H170 substitutions remodel and hydrate the pore, while H170D additionally creates a long-lived NH_4_^+^-binding trap.

## Discussion

The data presented here establish that NeRh50 uses the conserved Amt/Mep/Rh structural scaffold to support a transport mechanism that is fundamentally distinct from canonical AmtB^6–10^. More broadly, NeRh50 illustrates how an ancient transporter architecture can be evolutionarily repurposed into mechanistically distinct solutions adapted to different ecological niches, physiological roles and energetic constraints.

WT NeRh50 supports inward NH_4_^+^ uptake, but this activity contains two genetically and mechanistically separable electrogenic components: a D_2_O-sensitive pathway and a D_2_O-resistant pathway. In parallel, NeRh50 confers MeA protection in yeast, consistent with an export function. These three outputs are genetically dissociable: D162A retains export capacity but loses productive inward uptake, whereas H170 variants retain NH_4_^+^ translocation but lose both the D_2_O-resistant component and MeA protection. The mutant that loses the D_2_O-resistant pathway invariably loses export capacity with it. NeRh50 is therefore not an AmtB-like importer with an Rh label, it is a mechanistically distinct protein that has retained the ancestral import function while evolving an additional Rh-specific transport function.

Both transport components are best understood against the backdrop of the conserved Amt/Mep/Rh pore chemistry^6–8^. The WT NeRh50 energy landscape retains the dominant thermodynamic bias toward NH_3_ passage: NH_4_^+^ is stabilised in the periplasmic recruitment region but becomes strongly disfavoured as it enters the dehydrating gate, and the position-dependent pK_a_ profile places the NH_4_^+^/NH_3_ transition at the Phe-gate constriction, before the substrate reaches the twin-His motif. The Phe-gate therefore provides the dehydrating environment that favours deprotonation, whereas the twin-His motif functions as a downstream scaffold for proton and charge handling rather than as the primary deprotonation site. This D_2_O-sensitive component is mechanistically equivalent to the pathway established for AmtB and related transporters, in which NH_4_ is recruited at the periplasmic vestibule, deprotonation is favoured at the hydrophobic gate, and NH_3_ passage is coupled to proton movement through a solvent-exchangeable water network^19,20,44^.

The key difference is that NeRh50 carries out this conserved chemistry in a Rh-type pore that is structurally less stringent than AmtB^16^, and it is precisely this relaxation that creates the conditions for a D₂O-resistant pathway to coexist alongside the deprotonation-coupled route. In AmtB, the F107/F215 Phe-gate forms a tightly stacked aromatic barrier leading into a long, dehydrated hydrophobic pore exceeding 20 Å, making deprotonation at the gate tightly coupled to obligatory downstream proton transfer through the twin-His/water network^9,10,20^. In NeRh50, our simulations show that the F110/F218 gate is semi-open and asymmetric, with a mean inter-residue distance of 6.83 ± 1.44 Å. The effective hydrophobic barrier is also shorter, spanning only ∼11 Å and the twin-His region remains partially hydrated. These features do not abolish the NH_3_-favoured thermodynamic bias, but they reduce the structural penalty for transient accommodation of a charge-retaining NH_x_ species. In this architectural context, the mechanism proposed by Maeda *et al*. for AmtB becomes highly relevant in NeRh50: driven by the electrochemical gradient, equivalent in the SSME experiment to a ∼130-150 mV Nernst potential, NH_4_^+^ can transit in the pore without obligatory proton transfer through water wires^24^. Our unrestrained NH_4_^+^ simulations provide a structural context consistent with this model. In WT NeRh50, NH_4_^+^ interactions with the pore were transient, with no long-lived binding site detected within the central pathway. This dynamic behaviour is consistent with a pore that does not stably retain the charged substrate and remains compatible with transient passage of intact NH_4_^+^ under an appropriate electrochemical gradient. Because late cytoplasmic-face density arose from periodic-boundary crossing through bulk solvent, these trajectories were not used to assign individual translocation events. Instead, they support the view that the WT pore can transiently accommodate NH_4_^+^ without forming a stable central-pore intermediate.

The MeA-protective output is most simply explained as gradient-driven reversal of the same Rh-type electrogenic pathway. Transport through NeRh50 is not intrinsically directional; its direction is set by the prevailing electrochemical gradient. During ammonium acquisition the gradient favours inward transit; when Mep transporters drive MeA accumulation internally, the chemical gradient reverses and favours outward release through NeRh50. The semi-open Phe gate, shortened hydrophobic barrier and partially hydrated mid-pore collectively lower the energetic cost of outward substrate release, a structural permissiveness for reversal that the occluded AmtB gate does not allow, and that is entirely consistent with the bidirectional ammonium handling demonstrated for mammalian RhAG and RhCG^13,17,18^.

The mutagenesis data assign D162 and H170 to distinct arms of this branched transport landscape. D162 defines the entry-coupling arm: it links substrate recognition in the external vestibule to the canonical D_2_O-sensitive import pathway, and its loss reduces transport to a local binding event without productive translocation. H170 defines the Rh-specific arm: its imidazole chemistry maintains the pore in a state permissive to transient charge-retaining transit, and its loss, regardless of whether the substitution dehydrates the gate (H170A) or deepens the electrostatic interaction with NH_4_^+^ (H170D/E), abolishes both the D_2_O-resistant component and export capacity. That three chemically distinct substitutions with structurally opposite effects on gate geometry converge on the same functional loss identifies the imidazole ring chemistry itself, not pore hydration state, as the critical variable.

The Amt/Mep/Rh scaffold therefore operates not as a single fixed mechanism, but as a tuneable platform whose output is determined by local residue chemistry at a small number of pore positions. In NeRh50, this tuning creates a division of labour: selective uptake is preserved, but the pore also retains enough energetic flexibility to relieve intracellular nitrogen and charge stress when substrate accumulates. This is the adaptive value of the dual mechanism: NeRh50 can acquire ammonium without becoming a nonspecific cation channel, while still behaving as a reversible Rh-type valve when the physiological gradient changes.

This plasticity appears to have deep evolutionary roots. Phylogenomic evidence suggests that the last universal common ancestor (LUCA) already possessed genes for ammonium uptake, implying that ammonium transport was among the earliest biological solutions to nitrogen acquisition, predating complete biological nitrogen fixation^49^. The later diversification of the family into importers, sensors and bidirectional Rh-type transporters can be viewed as evolutionary retuning of an ancient pore under shifting ecological pressure: scarcity favoured stringent acquisition, abundance favoured detoxifying release, and fluctuating environments favoured sensing. This reframes how the Amt/Mep/Rh superfamily should be understood: not as a set of phylogenetically distinct transporter classes, each with its own fixed mechanism, but as a single conserved scaffold whose pore landmarks function as evolutionary control points featuring residue-level substitutions that modify directionality, coupling and physiological output without altering the underlying fold. Ammonium transport has evolved not through the emergence of a new architecture, but through repeated repurposing of an existing scaffold.

## Supporting information

Molecular dynamics supplementary data

## Notes

### Competing Interest Statement

The authors have declared no competing interest.

## References

1. Gerlt, J. A. and Babbitt, P. C. (2001). Divergent evolution of enzymatic function: mechanistically diverse superfamilies and functionally distinct suprafamilies. Annual Review of Biochemistry. doi:10.1146/annurev.biochem.70.1.209

2. Akiva, E., Copp, J. N., Tokuriki, N. and Babbitt, P. C. (2017). Evolutionary and molecular foundations of multiple contemporary functions of the nitroreductase superfamily. Proceedings of the National Academy of Sciences USA. doi:10.1073/pnas.1706849114

3. Babst, M. (2020). Regulation of nutrient transporters by metabolic and environmental stresses. Current Opinion in Cell Biology. doi:10.1016/j.ceb.2020.02.009

4. Darbani, B., Kell, D. B. and Borodina, I. (2018). Energetic evolution of cellular transportomes. BMC Genomics. doi:10.1186/s12864-018-4816-5

5. Karapanagioti, F., Atlason, Ú. Á., Slotboom, D. J., Poolman, B. and Obermaier, S. (2024). Fitness landscape of substrate-adaptive mutations in evolved amino acid-polyamine-organocation transporters. eLife. doi:10.7554/eLife.93971

6. Bizior, A., Williamson, G., Harris, T., Hoskisson, P. A. and Javelle, A. (2023). Prokaryotic ammonium transporters: what has three decades of research revealed? Microbiology. doi:10.1099/mic.0.001360

7. Williamson, G., Bizior, A., Harris, T., Pritchard, L., Hoskisson, P. A. and Javelle, A. (2024). Biological ammonium transporters from the Amt/Mep/Rh superfamily: mechanism, energetics, and technical limitations. Bioscience Reports. doi:10.1042/BSR20211209

8. Williamson, G., Harris, T., Bizior, A., Hoskisson, P. A., Pritchard, L. and Javelle, A. (2024). Biological ammonium transporters: evolution and diversification. The FEBS Journal. doi:10.1111/febs.17059

9. Khademi, S., O’Connell, J., Remis, J., Robles-Colmenares, Y., Miercke, L. J. W. and Stroud, R. M. (2004). Mechanism of ammonia transport by Amt/MEP/Rh: structure of AmtB at 1.35 Å. Science. doi:10.1126/science.1101952

10. Zheng, L., Kostrewa, D., Bernèche, S., Winkler, F. K. and Li, X.-D. (2004). The mechanism of ammonia transport based on the crystal structure of AmtB of Escherichia coli. Proceedings of the National Academy of Sciences USA. doi:10.1073/pnas.0406475101

11. Andrade, S. L. A., Dickmanns, A., Ficner, R. and Einsle, O. (2005). Crystal structure of the archaeal ammonium transporter Amt-1 from Archaeoglobus fulgidus. Proceedings of the National Academy of Sciences USA. doi:10.1073/pnas.0506208102

12. van den Berg, B., Chembath, A., Jefferies, D., Basle, A., Khalid, S. and Rutherford, J. C. (2016). Structural basis for Mep2 ammonium transceptor activation by phosphorylation. Nature Communications. doi:10.1038/ncomms11337

13. Gruswitz, F., Chaudhary, S., Ho, J. D., Schlessinger, A., Pezeshki, B., Ho, C.-M., Sali, A., Westhoff, C. M. and Stroud, R. M. (2010). Function of human Rh based on structure of RhCG at 2.1 Å. Proceedings of the National Academy of Sciences USA. doi:10.1073/pnas.1003587107

14. Lupo, D., Li, X.-D., Durand, A., Tomizaki, T., Cherif-Zahar, B., Matassi, G., Merrick, M. and Winkler, F. K. (2007). The 1.3-Å resolution structure of Nitrosomonas europaea Rh50 and mechanistic implications for NH3 transport by Rhesus family proteins. Proceedings of the National Academy of Sciences USA. doi:10.1073/pnas.0706563104

15. Li, X., Jayachandran, S., Nguyen, H.-H. T. and Chan, M. K. (2007). Structure of the Nitrosomonas europaea Rh protein. Proceedings of the National Academy of Sciences USA. doi:10.1073/pnas.0709710104

16. Cooper, B., Bizior, A., Talandashti, R., Oluwole, A., Henderson, P., Harris, T., Robinson, C., Hoskisson, P. A., Khalid, S., Pritchard, L., Isom, G. and Javelle, A. (2026). Lipid-associated architecture and divergent ammonium transport distinguish a bacterial Rhesus protein. bioRxiv. doi:10.64898/2026.09.17.752382

17. Marini, A. M., Matassi, G., Raynal, V., André, B., Cartron, J.-P. and Cherif-Zahar, B. (2000). The human Rhesus-associated RhAG protein and a kidney homologue promote ammonium transport in yeast. Nature Genetics. doi:10.1038/81656

18. Zidi-Yahiaoui, N., Mouro-Chanteloup, I., D’Ambrosio, A.-M., Lopez, C., Gane, P., Le Van Kim, C., Cartron, J.-P., Colin, Y. and Ripoche, P. (2005). Human Rhesus B and Rhesus C glycoproteins: properties of facilitated ammonium transport in recombinant kidney cells. Biochemical Journal. doi:10.1042/BJ20050657

19. Javelle, A., Lupo, D., Ripoche, P., Fulford, T., Merrick, M. and Winkler, F. K. (2008). Substrate binding, deprotonation, and selectivity at the periplasmic entrance of the Escherichia coli ammonia channel AmtB. Proceedings of the National Academy of Sciences USA. doi:10.1073/pnas.0711742105

20. Williamson, G., Tamburrino, G., Bizior, A., Boeckstaens, M., Dias Mirandela, G., Bage, M. G., Pisliakov, A., Ives, C. M., Terras, E., Hoskisson, P. A., Marini, A. M., Zachariae, U. and Javelle, A. (2020). A two-lane mechanism for selective biological ammonium transport. eLife. doi:10.7554/eLife.57183

21. Mak, D. O. D., Dang, B., Weiner, I. D., Foskett, J. K. and Westhoff, C. M. (2006). Characterization of ammonia transport by the kidney Rh glycoproteins RhBG and RhCG. American Journal of Physiology-Renal Physiology. doi:10.1152/ajprenal.00147.2005

22. Dias Mirandela, G., Tamburrino, G., Ivanović, M. T., Strnad, F. M., Byron, O., Rasmussen, T., Hoskisson, P. A., Hub, J. S., Zachariae, U., Gabel, F. and Javelle, A. (2018). Merging in-solution X-ray and neutron scattering data allows fine structural analysis of membrane-protein detergent complexes. Journal of Physical Chemistry Letters. doi:10.1021/acs.jpclett.8b01598

23. Wang, S., Orabi, E. A., Baday, S., Bernèche, S. and Lamoureux, G. (2012). Ammonium transporters achieve charge transfer by fragmenting their substrate. Journal of the American Chemical Society. doi:10.1021/ja300129x

24. Maeda, K., Kurata, H., Javelle, A., Westerhoff, H. V. and Boogerd, F. C. (2026). Computer experimentation reveals mechanisms for signaling and coupled transport that trade efficiency for robust growth. npj Systems Biology and Applications. doi:10.1038/s41540-026-00793-1

25. Chain, P., Lamerdin, J., Larimer, F., Regala, W., Lao, V., Land, M., Hauser, L., Hooper, A., Klotz, M., Norton, J., Sayavedra-Soto, L., Arciero, D., Hommes, N., Whittaker, M. and Arp, D. (2003). Complete genome sequence of the ammonia-oxidizing bacterium and obligate chemolithoautotroph Nitrosomonas europaea. Journal of Bacteriology. doi:10.1128/JB.185.9.2759-2773.2003

26. Arp, D. J., Sayavedra-Soto, L. A. and Hommes, N. G. (2002). Molecular biology and biochemistry of ammonia oxidation by Nitrosomonas europaea. Archives of Microbiology. doi:10.1007/s00203-002-0452-0

27. Cherif-Zahar, B., Durand, A., Schmidt, I., Hamdaoui, N., Matic, I., Merrick, M. and Matassi, G. (2007). Evolution and functional characterization of the RH50 gene from the ammonia-oxidizing bacterium Nitrosomonas europaea. Journal of Bacteriology. doi:10.1128/JB.01089-07

28. Coutts, G., Thomas, G., Blakey, D. and Merrick, M. (2002). Membrane sequestration of the signal transduction protein GlnK by the ammonium transporter AmtB. The EMBO Journal. doi:10.1093/emboj/21.4.536

29. Rentsch, D., Laloi, M., Rouhara, I., Schmelzer, E., Delrot, S. and Frommer, W. B. (1995). NTR1 encodes a high-affinity oligopeptide transporter in Arabidopsis. FEBS Letters. doi:10.1016/0014-5793(95)00853-2

30. Marini, A. M., Soussi-Boudekou, S., Vissers, S. and André, B. (1997). A family of ammonium transporters in Saccharomyces cerevisiae. Molecular and Cellular Biology. doi:10.1128/MCB.17.8.4282

31. Williamson, G., Brito, A. S., Bizior, A., Tamburrino, G., Dias Mirandela, G., Hoskisson, P. A., Marini, A. M., Zachariae, U. and Javelle, A. (2022). Coexistence of ammonium transporter and channel mechanisms in Amt-Mep-Rh twin-His variants impairs the filamentation signaling capacity of fungal Mep2 transceptors. mBio. doi:10.1128/mbio.02913-21

32. Mirandela, G. D., Tamburrino, G., Hoskisson, P. A., Zachariae, U. and Javelle, A. (2019). The lipid environment determines the activity of the Escherichia coli ammonium transporter AmtB. The FASEB Journal. doi:10.1096/fj.201800782R

33. R Core Team. (2025). R: A language and environment for statistical computing. R Foundation for Statistical Computing, Vienna, Austria.

34. Lüdecke, D., Ben-Shachar, M. S., Patil, I. and Makowski, D. (2020). Extracting, computing and exploring the parameters of statistical models using R. Journal of Open Source Software 5, 2445. doi:10.21105/joss.02445

35. Lüdecke, D., Ben-Shachar, M. S., Patil, I., Waggoner, P. and Makowski, D. (2021). performance: An R package for assessment, comparison and testing of statistical models. Journal of Open Source Software 6, 3139. doi:10.21105/joss.03139

36. Lenth, R. V. and Piaskowski, J. (2026). emmeans: Estimated Marginal Means, aka Least-Squares Means. R package version 2.0.2. doi:10.32614/CRAN.package.emmeans

37. GraphPad Software. (2023). GraphPad Prism version 11.0.0 for Windows. GraphPad Software, Boston, Massachusetts USA. www.graphpad.com

38. Lee, J., Patel, D. S., Ståhle, J., Park, S.-J., Kern, N. R., Kim, S., Lee, J., Cheng, X., Valvano, M. A., Holst, O., Knirel, Y. A., Qi, Y., Jo, S., Klauda, J. B., Widmalm, G. and Im, W. (2019). CHARMM-GUI Membrane Builder for complex biological membrane simulations with glycolipids and lipoglycans. Journal of Chemical Theory and Computation. doi:10.1021/acs.jctc.8b01066

39. Huang, J., Rauscher, S., Nawrocki, G., Ran, T., Feig, M., de Groot, B. L., Grubmüller, H. and MacKerell, A. D. Jr (2017). CHARMM36m: an improved force field for folded and intrinsically disordered proteins. Nature Methods. doi:10.1038/nmeth.4067

40. van der Spoel, D., Lindahl, E., Hess, B., Groenhof, G., Mark, A. E. and Berendsen, H. J. C. (2005). GROMACS: fast, flexible, and free. Journal of Computational Chemistry. doi:10.1002/jcc.20291

41. Hub, J. S., de Groot, B. L. and van der Spoel, D. (2010). g_wham—A free weighted histogram analysis implementation including robust error and autocorrelation estimates. Journal of Chemical Theory and Computation. doi:10.1021/ct100494z

42. Michaud-Agrawal, N., Denning, E. J., Woolf, T. B. and Beckstein, O. (2011). MDAnalysis: a toolkit for the analysis of molecular dynamics simulations. Journal of Computational Chemistry. doi:10.1002/jcc.21787

43. Bates, R. G. and Pinching, G. D. (1949). Acidic dissociation constant of ammonium ion at 0° to 50 °C, and the base strength of ammonia. Journal of Research of the National Bureau of Standards. doi:10.6028/jres.042.037

44. Ariz, I., Boeckstaens, M., Gouveia, C., Martins, A. P., Sanz-Luque, E., Fernández, E., Soveral, G., von Wirén, N., Marini, A. M., Aparicio-Tejo, P. M. and Cruz, C. (2018). Nitrogen isotope signature evidences ammonium deprotonation as a common transport mechanism for the AMT-Mep-Rh protein superfamily. Science Advances. doi:10.1126/sciadv.aar3599

45. Bazzone, A., Barthmes, M. and Fendler, K. (2017). SSM-based electrophysiology for transporter research. Methods in Enzymology. doi:10.1016/bs.mie.2017.05.008

46. Deschuyteneer, G., El Bakkoury, M., Peulen, O., Cambier, P., Forêt, M., Laduron, S., Marini, A. M. and Boeckstaens, M. (2013). Rh proteins can mediate methylammonium export in yeast. PLoS ONE. doi:10.1371/journal.pone.0071092

47. Akgun, U. and Khademi, S. (2011). Periplasmic vestibule plays an important role for solute recruitment, selectivity, and gating in the Rh/Amt/MEP superfamily. Proceedings of the National Academy of Sciences USA. doi:10.1073/pnas.1007240108

48. Nygaard, T. P., Alfonso-Prieto, M., Peters, G. H., Jensen, M. Ø. and Rovira, C. (2010). Substrate recognition in the Escherichia coli ammonia channel AmtB: a QM/MM investigation. Journal of Physical Chemistry B. doi:10.1021/jp102338h

49. Boden, J. S., Ni, Z., Anderson, R. E. and Stüeken, E. E. (2026). Abiotic sources of fixed nitrogen sustained early ecosystems for several hundred million years after the origin of life. Science Advances, 12(25), eaec4450. doi:10.1126/sciadv.aec4450

50. Seiferth, D. and Biggin, P. C. (2024). Exploring the influence of pore shape on conductance and permeation. Biophysical Journal, 123(18), 3107–3119. doi:10.1016/j.bpj.2024.07.010

