## Supplementary material for "A bacterial Rhesus transporter retunes a structurally conserved ammonium pore into a reversible nitrogen valve": Molecular dynamics supplementary data

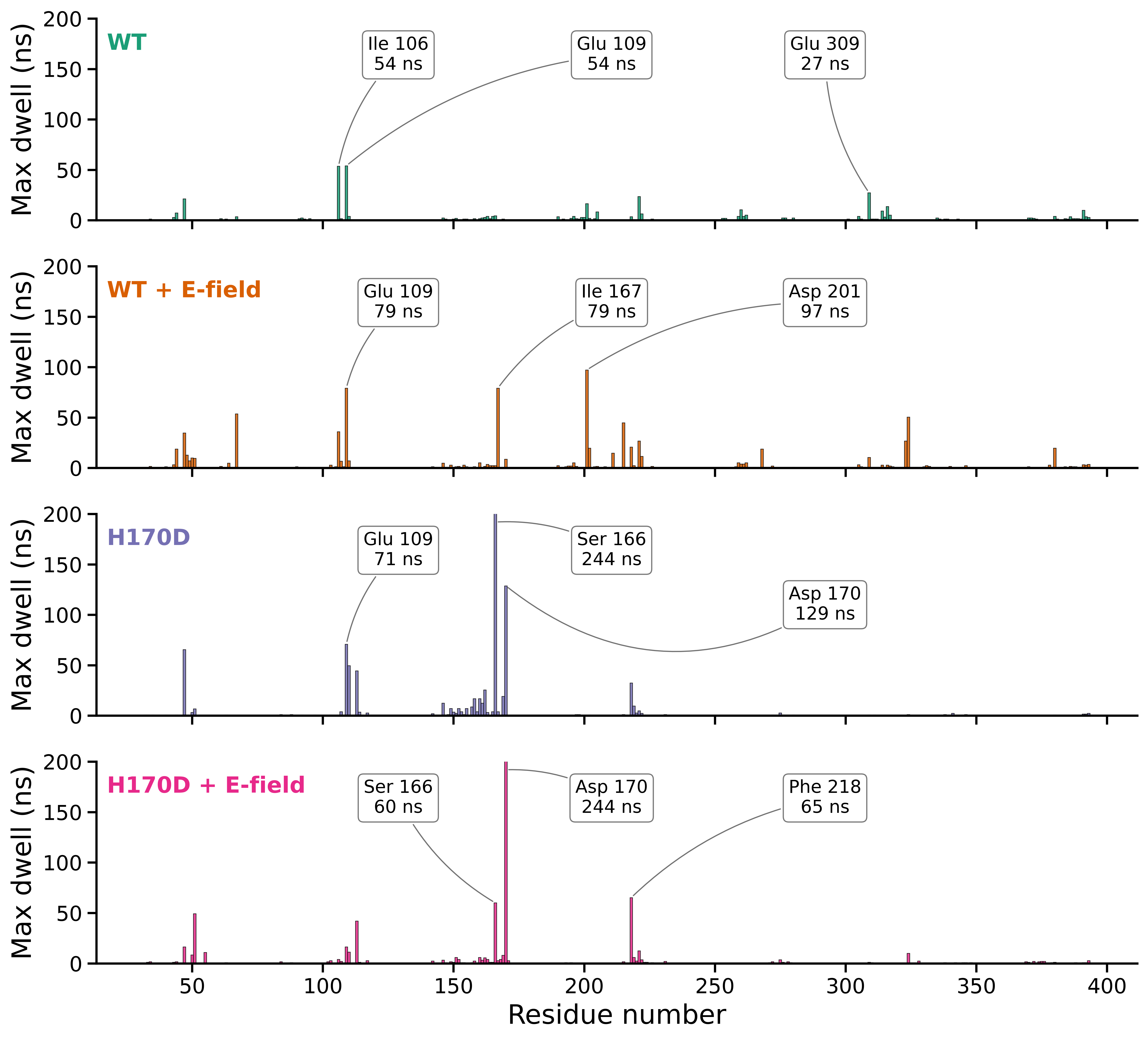

***Supplementary Figure S1. H170D creates a long-lived NH_4_^+^-retention site at the upper gate*** *Per-residue maximum NH_4_^+^ dwell times were calculated for WT and H170D NeRh50 in simulations performed with or without a constant electric field (0.05 V nm^-1^ along −z). Three NH_4_^+^ ions were initially placed in the periplasm-facing vestibule, one per protomer, and followed in three independent 250-ns trajectories per condition. For each residue, bars show the longest continuous contact observed across the trajectories using the contact criterion defined in Methods. Selected maximum dwell times are annotated. In WT, dwell events were comparatively short and distributed across several residues. In H170D, long-lived occupancy was concentrated around S166 and the introduced D170 residue. Under the applied field, the maximum dwell time at D170 reached 244 ns, approaching the full trajectory duration. These maximum dwell times quantify kinetic retention rather than equilibrium binding affinity and do not demonstrate complete NH_4_^+^ permeation through the pore.*

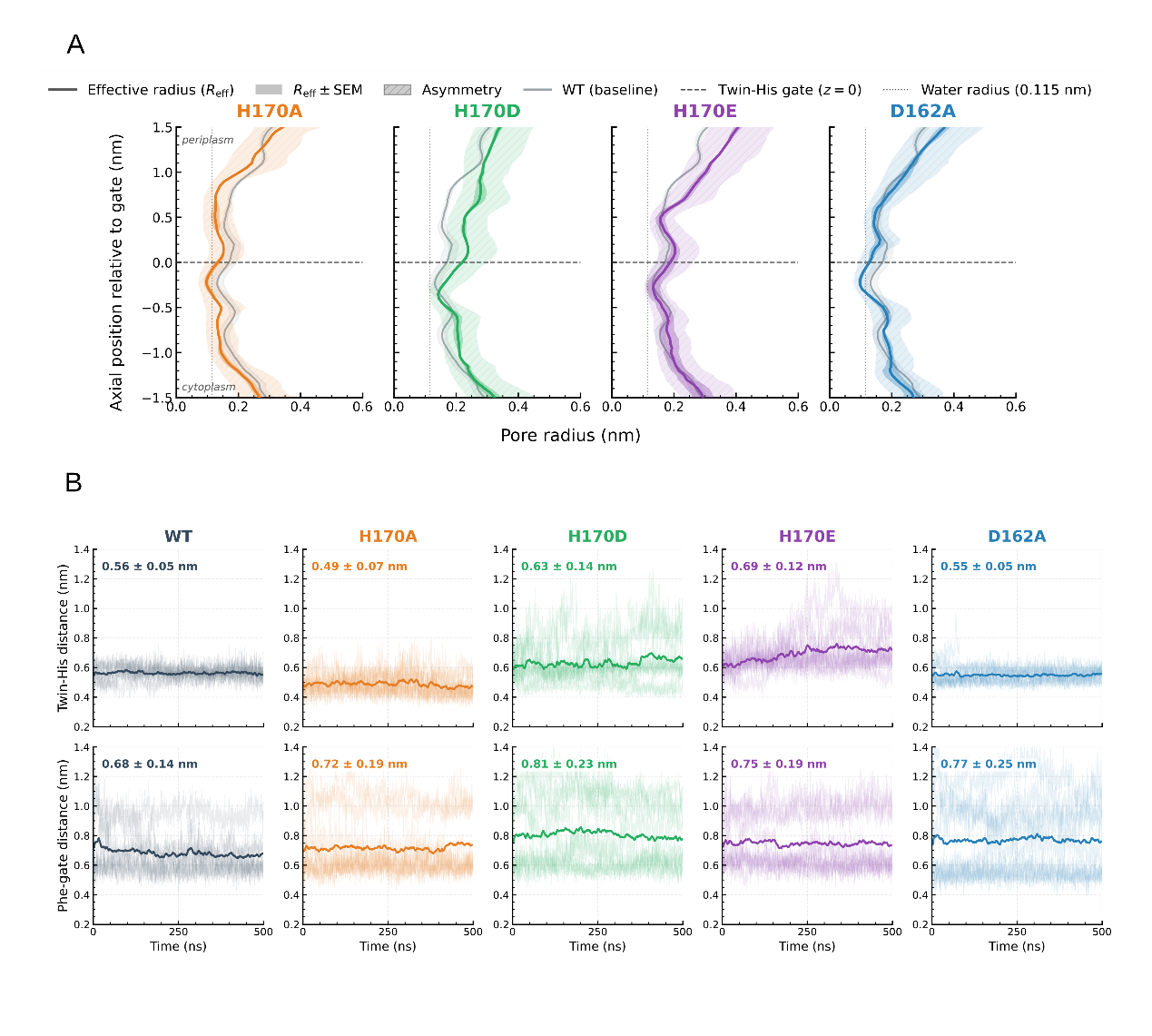
***Supplementary Figure S2. H170 substitutions remodel the pore dimensions and gate geometry of NeRh50.***

*Analyses were performed using three independent 500-ns substrate-free simulations of WT NeRh50 and the indicated variants.*

***(A)*** *Axial effective pore-radius profiles for H170A, H170D, H170E and D162A, calculated using PoreAnalyser*^50^, each compared with the WT baseline shown in grey. The horizontal dashed line marks the twin-His gate at *Z=0, and the vertical dotted line indicates the radius of a water molecule (0.115 nm). Positive and negative axial coordinates correspond to the periplasm- and cytoplasm-facing sides, respectively.*

***(B)*** *Time evolution of the distance between residues 170 and 324 (top) and the Phe-gate distance (bottom). For the H170 variants, the upper measurement represents the distance between the substituted residue at position 170 and H324. Faint traces show the underlying trajectory data, whereas bold curves show the corresponding averaged trends. H170D and H170E increased both the 170-324 separation and the Phe-gate distance relative to WT, consistent with dilation of the central constrictions. H170A reduced the 170–324 separation, whereas D162A retained a WT-like 170-324 distance but increased the Phe-gate separation. Together, these analyses show that the substitutions remodel the two gates in mutation-specific rather than uniformly opening directions.*
